# Structures of core eukaryotic protein complexes

**DOI:** 10.1101/2021.09.30.462231

**Authors:** Ian R. Humphreys, Jimin Pei, Minkyung Baek, Aditya Krishnakumar, Ivan Anishchenko, Sergey Ovchinnikov, Jing Zhang, Travis J. Ness, Sudeep Banjade, Saket Bagde, Viktoriya G. Stancheva, Xiao-Han Li, Kaixian Liu, Zhi Zheng, Daniel J. Barrero, Upasana Roy, Israel S. Fernández, Barnabas Szakal, Dana Branzei, Eric C. Greene, Sue Biggins, Scott Keeney, Elizabeth A. Miller, J. Christopher Fromme, Tamara L. Hendrickson, Qian Cong, David Baker

**Affiliations:** Department of Biochemistry, University of Washington, Seattle, WA, USA; Institute for Protein Design, University of Washington, Seattle, WA, USA; Eugene McDermott Center for Human Growth and Development, University of Texas Southwestern Medical Center, Dallas, TX, USA; Department of Biophysics, University of Texas Southwestern Medical Center, Dallas, TX, USA; Faculty of Arts and Sciences, Division of Science, Harvard University, Cambridge, MA 02138, USA; John Harvard Distinguished Science Fellowship Program, Harvard University, Cambridge, MA, USA; Department of Chemistry, Wayne State University, Detroit, MI, USA; Department of Molecular Biology & Genetics, Weill Institute for Cell and Molecular Biology, Cornell University, Ithaca, NY, USA; MRC Laboratory of Molecular Biology, Cambridge, CB2 0QH, UK; Molecular Biology Program, Memorial Sloan Kettering Cancer Center, New York, NY; Gerstner Sloan Kettering Graduate School of Biomedical Sciences, New York, NY; Howard Hughes Medical Institute, Division of Basic Sciences, Fred Hutchinson Cancer Research Center, 1100 Fairview Avenue N, Seattle, WA, USA; Department of Biochemistry and Molecular Biophysics, Columbia University, New York, NY, USA; Department of Structural Biology, St Jude Children’s Research Hospital, Memphis, TN, USA; IFOM, the FIRC Institute of Molecular Oncology, Via Adamello 16, 20139, Milan, Italy; Istituto di Genetica Molecolare, Consiglio Nazionale delle Ricerche (IGM-CNR), Via Abbiategrasso 207, 27100, Pavia, Italy; Howard Hughes Medical Institute, Memorial Sloan Kettering Cancer Center, New York, NY, USA; Howard Hughes Medical Institute, University of Washington, Seattle, WA, USA

**Author notes:** Contributed equally (Ian R. Humphreys, Jimin Pei, Minkyung Baek, Aditya Krishnakumar). Contributed equally (Qian Cong, David Baker).

## Abstract

Protein-protein interactions play critical roles in biology, but despite decades of effort, the structures of many eukaryotic protein complexes are unknown, and there are likely many interactions that have not yet been identified. Here, we take advantage of recent advances in proteome-wide amino acid coevolution analysis and deep-learning-based structure modeling to systematically identify and build accurate models of core eukaryotic protein complexes, as represented within the *Saccharomyces cerevisiae* proteome. We use a combination of RoseTTAFold and AlphaFold to screen through paired multiple sequence alignments for 8.3 million pairs of *S. cerevisiae* proteins and build models for strongly predicted protein assemblies with two to five components. Comparison to existing interaction and structural data suggests that these predictions are likely to be quite accurate. We provide structure models spanning almost all key processes in Eukaryotic cells for 104 protein assemblies which have not been previously identified, and 608 which have not been structurally characterized.

**One-sentence summary:** We take advantage of recent advances in proteome-wide amino acid coevolution analysis and deep-learning-based structure modeling to systematically identify and build accurate models of core eukaryotic protein complexes.

---

Yeast two hybrid (Y2H), affinity-purification mass spectrometry (APMS), and other high-throughput experimental approaches have identified many pairs of interacting proteins in yeast and other organisms (*1*)(*2*)(*3*)(*4*)(*5*), but there are often extensive discrepancies between sets generated using the different methods and considerable false positive and false negative rates (*6*). Since residues at protein-protein interfaces are expected to coevolve, given two proteins, the likelihood that they interact can be assessed by identifying and aligning the sequences of orthologs of the two proteins in many different species, joining them to create paired multiple sequence alignments (pMSA), and then determining the extent to which changes in the sequences of orthologs for the first partner covary with ortholog sequence changes for the second partner (*7*)(*8*). Such amino acid coevolution has been used to guide modeling of complexes for cases in which the structures of the partners are known (*9*)(*10*), and to systematically identify pairs of interacting proteins in Prokaryotes with accuracy higher than experimental screens (*7*). Recent deep-learning-based advances in protein structure prediction have the potential to increase the power of such approaches as they (*11*)(*12*) now enable accurate modeling not only of protein monomer structures but also protein complexes (*11*).

We set out to combine proteome wide coevolution-guided protein interaction identification with deep learning based protein structure modeling to systematically identify and determine the structures of eukaryotic protein assemblies. We faced several challenges in directly applying to eukaryotes the statistical methods effective in identifying coevolving pairs in prokaryotes. First, far more genome sequences are available for prokaryotes than eukaryotes, and the average number of homologous amino acid sequences (excluding nearly identical copies with > 95% sequence identity) is on the order of 10,000 for bacterial proteins, but 1,000 for eukaryotic proteins (fig. S1). Thus, multiple sequence alignments for pairs of eukaryotic proteins contain far fewer diverse sequences, making it more difficult for statistical methods to distinguish true coevolutionary signal from the noise. Second, eukaryotes in general have a larger number of genes, making comprehensive pairwise analysis more computationally intensive, and increasing the background noise resulting from calculation errors. Third, mRNA splicing in eukaryotes further increases the number of protein species, resulting in errors in gene predictions and complicating sequence alignments. Fourth, eukaryotes underwent several rounds of genome duplications in multiple lineages (*13*), and it can be difficult to distinguish orthologs from paralogs, which is important for detecting coevolutionary signal because the protein interactions of interest are likely to be conserved in orthologs in other species but less so in paralogs.

We sought to overcome these challenges as follows. To help with the first three challenges, we chose the yeast *S. cerevisiae* as the starting point because there are a large number (~1,700) of fungal genomes (*14*), the genome is relatively small (6,000 genes total), and there is relatively little mRNA splicing (*15*). Furthermore, because the interactome of yeast has been extensively studied, there is a “gold standard” set of known interactions to evaluate the reliability of predicted interactions and structures.

To distinguish orthologs from paralogs, we started from OrthoDB (*16*), a hierarchical catalog of orthologs over 1,271 Eukaryote genomes, and supplemented each orthologous group with sequences from 4,325 Eukaryote proteomes we assembled from NCBI (https://www.ncbi.nlm.nih.gov/genome) and JGI (*17*). We compared the protein sequences for each of the additional 4,325 proteomes against those of the most closely related species in the OrthoDB database, and used the reciprocal best hit criterion (*18*) to identify orthologs; these were then added to the corresponding orthologous group. A complication is that each species frequently contains multiple proteins belonging to the same orthologous group, leading to ambiguity in determining which protein should be included in pMSAs crucial for coevolutionary analysis. These multiple copies may represent alternatively spliced forms of the same gene, parts of the same gene that were split into multiple pieces due to errors in gene prediction, or recent gene expansions specific to certain lineages. We dealt with these possibilities by keeping only the longest isoform of each gene, merging pieces of the same gene, and selecting the copy with the highest sequence identity to single-copy orthologs in other species. For 4,090 out of ~6,000 yeast proteins, we were able to identify clear orthologs across large numbers of species, and we generated pMSAs for all 4,090 * 4,089 / 2 = 8,362,005 pairwise combinations of these proteins. We focused on 4,286,433 pairs with alignments containing over 200 sequences to increase prediction accuracy and less than 1,300 amino acids to allow fast computation.

In a first set of calculations, we found that even with the advantages of *S. cerevisiae* and improved ortholog identification, the statistical method (Direct Coupling Analysis, DCA) we had used in our previous coevolution-guided PPI screen in Prokaryotes (*7*) (the more accurate GREMLIN (*9*) method is too slow for this) could not effectively distinguish a “gold standard” set of 768 yeast protein pairs known to interact (*5*) (http://interactome.dfci.harvard.edu/S_cerevisiae/) from the much larger set (768,000 pairs) of primarily non-interacting pairs (Fig. 1A, grey curve, area under the curve: 0.016). Progress clearly required a more accurate and sensitive, but still rapidly computable, method for evaluating protein interactions based on pMSAs.

**Figure 1.**
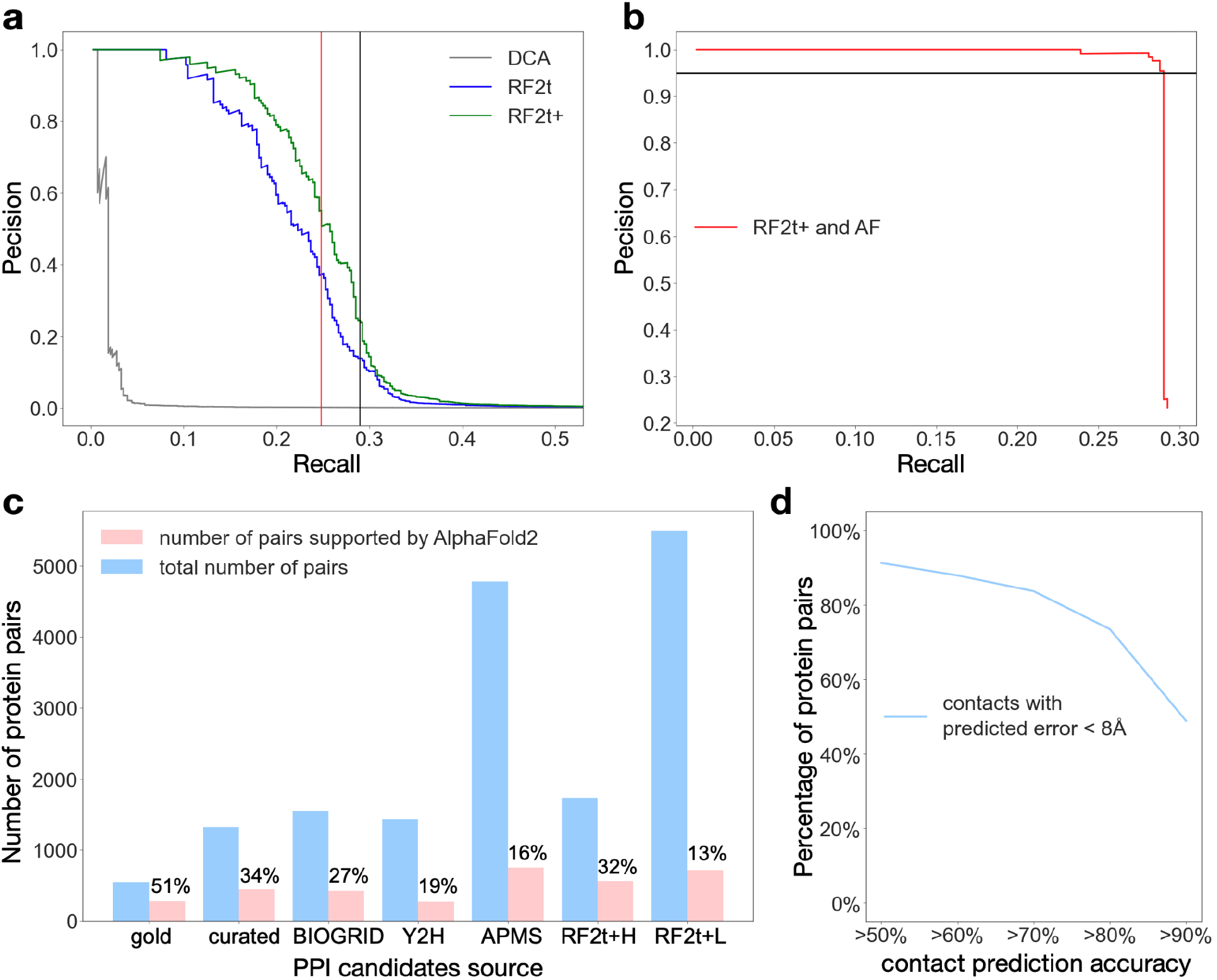
Benchmark of *in silico* PPI screen and protein complex modeling. (**a**) Performance (precision at different levels of recall) of different methods in picking out gold standard interacting pairs from the set of 4.3 million pMSAs. Pairs were ranked by the top coevolution score or contact probability between residue pairs. DCA: Direct coupling analysis. RF2t: top contact probability between residues of two proteins by RF 2-track model. RF2t+, optimized RF2t with two modifications: (1) the C-terminal residues (10 amino acids) of the first protein and the N-terminal residues (10 amino acids) of the second protein was excluded when computing the top inter-residue contact probability for a protein pair; (2) average product correction was applied to the contact probability of a protein pair to penalize proteins that have high contact probability with many other proteins. (**b**) Performance (precision at different levels of recall) in picking out gold standard protein interaction amongst the 5495 protein pairs selected by the RF2t+ method in panel a. Three models (1, 3, and 5) were generated by AlphaFold, and the top residue-residue contact probability excluding the 10 residues near the junction between two proteins were calculated for each model. The minimum top contact probability among the 3 models was used to indicate the probability for two proteins to interact. (**c**) Application of AlphaFold to screen PPI candidates from different sources. (**d**) Distribution of percent of inter-protein contacts predicted by AF that are observed as contacts (< 8Å) in closely-related experimental structures.

We explored the application of the recently developed deep learning based structure prediction methods, RoseTTAFold (RF) and AlphaFold (AF), to this problem. Even though RF was originally trained on monomeric protein sequences and structures, it can accurately predict the structures of protein complexes given pMSAs with a sufficient number of sequences (*11*). We evaluated the compute time required and the PPI prediction accuracy for a variety of model architectures, and found that a lighter-weight (10.7 million parameters) RF two-track model provided an optimal tradeoff: the model requires 11 seconds (about 100 times faster than AF) to process a pMSA of 1,000 amino acids on a NVIDIA TITAN RTX graphic processing unit, and it can effectively distinguish gold standard PPIs amongst much larger sets of randomly paired proteins. The very short time required to analyze an individual pMSA made it possible to process all 4.3 million pMSAs. This method considerably outperformed DCA in distinguishing gold standard interactions from random pairs (Fig. 1A, blue curve, area under the curve: 0.219), using the highest predicted contact probability over all pairs of residues in the two proteins as a measure of the propensity for two proteins to interact. Performance was further improved (Fig. 1A, green curve, area under the curve: 0.248) by correcting overestimations of predicted contact probabilities between the C-terminal residues of the first protein and the N-terminal residues of the second protein, and of predicted interactions for a subset of proteins showing hub-like interactions with many other proteins. The much better performance of RF than DCA likely stems from the extensive information on protein sequence-structure relationships embedded in the RF deep neural network; DCA by contrast operates solely on protein sequences with no underlying protein structure model.

We next explored whether AF residue-residue contact predictions could further distinguish interacting from non-interacting protein pairs. Like RF, AF was trained on monomeric protein structures, but given the good results with 2-track RF on protein complexes, and the higher accuracy of AF (also a 2-track network with a final 3D structure module) on monomers, we reasoned that it might similarly have higher accuracy on complexes. To enable modeling of protein complexes using AF, we modified the positional encoding. AF was too slow to be applied to the entire set of 4.3 million pMSAs (this would require 0.1-1 million GPU hours); instead we applied AF to the 5,495 protein pairs with the highest RF support (corresponding to ~25% precision and ~29% recall based on our benchmark, indicated by the black vertical line in Fig. 1A). Using the highest AF contact probability over all residue pairs as a measure of interaction strength, we found that the combination of RF followed by AF provided excellent performance (Fig. 1B). Almost all the gold-standard pairs were ranked higher than the negative controls by AF contact probability, allowing selection of a set of 717 candidate PPIs with an expected precision of 95% at an AF contact probability cutoff of 0.67 (black line in Fig. 1B); we refer to this RF plus AF procedure as the *de novo* PPI screen, and the resulting set of predicted interactions, the *de novo* PPI set, below.

Due to the tradeoff between compute time and accuracy, and the necessity of setting a stringent threshold to avoid large numbers of false positives given the very large number of total pairs, we were concerned that some interacting proteins might not coevolve sufficiently to be identified robustly in our all-vs-all RF screen. Given the excellent performance of AF in distinguishing gold standard interactions amongst the RF filtered pMSAs, we also applied AF to pMSAs for PPIs reported in literature, including those identified in experimental high throughput screens. Similarly to our *de novo* PPI screen procedure, we considered protein pairs with AF contact probability larger than 0.67 to be confident interacting partners. We found that 51% of the gold standard PPI was supported by high AF contact probability (Fig. 1C), with lower ratios for candidate PPIs manually curated from multiple literature (http://interactome.dfci.harvard.edu/S_cerevisiae/download/LC_multiple.txt) (*3*) (34%) or supported by low-throughput experiments according to BIOGRID (*19*) (27%). The ratio of AF-supported PPIs is even lower for protein pairs identified by Y2H (19%) or APMS (16%) screens, consistent with the known larger fraction of false positives in large-scale experimental screens (*20*). The fast RF 2-track model used in the *de novo* screen has comparable or better accuracy than the large-scale experimental screens when assessed in this way: with a high stringency RF cutoff, the fraction of AF-supported pairs among PPIs identified by RF is 32%, similar to the accuracy of low-throughput experiments; with a lower stringency cutoff, this fraction becomes closer to that of the large scale experimental screens but somewhat fewer true PPIs are missed (Fig 1C).

In total, we identified 717 likely interacting pairs from the “*de novo* RF → AF” screen, and 1,223 from the “pooled experimental sets → AF” screen, of which 434 overlap, resulting in a total of 1,506 PPIs. Out of these, 718 have been structurally characterized, 684 have some supporting experimental data from literature and databases, and 104 are not to our knowledge previously described. To evaluate the accuracy of the predicted 3D structure of protein complexes, we used as a benchmark the 718 pairs with experimental structure in PDB. For 90% of these pairs, the majority of AF-predicted contacts are present in the previously determined structures (Fig. 1D).

With these benchmark results providing confidence in the accuracy of the new complex interaction predictions and 3D models of the predicted complexes, we next proceeded to analyze the structure models for the 852 complexes for which high resolution structural information was not previously available. Structure models of these complexes can be downloaded at **xxx**. We classified these into groups based on their biological functions, and provide examples of complexes in each functional class in Figs. 2–4. A first set of complexes are involved in maintenance and processing of genetic information: DNA repair, mitosis and meiosis checkpoints, transcription, and translation (Fig. 2). A second set of complexes play roles in protein translocation, transport through the secretory pathway, the cytoskeleton, and in mitochondria (Fig. 3). A third set of complexes are involved in metabolism (Fig. 4A). Protein complexes in which proteins of unknown function are predicted to interact with well characterized ones are shown in Fig. 4B: these interactions provide hints about the function of the uncharacterized proteins and could help identify new components of previously characterized assemblies. In cases where three or more proteins were predicted to mutually interact, we generated models of the full assemblies by concatenating the multiple sequence alignments for all of the components as illustrated by examples in Fig. 5; in most such cases the pairwise interactions in such models are quite similar to those for the independently built binary complexes, but simultaneous modeling of the full complex has the advantage of allowing conformational changes that could accompany full assembly. We do not have space to fully describe each of these new complexes, instead we focus on a few examples to illustrate the biological insights that can be drawn from these models.

**Figure 2.**
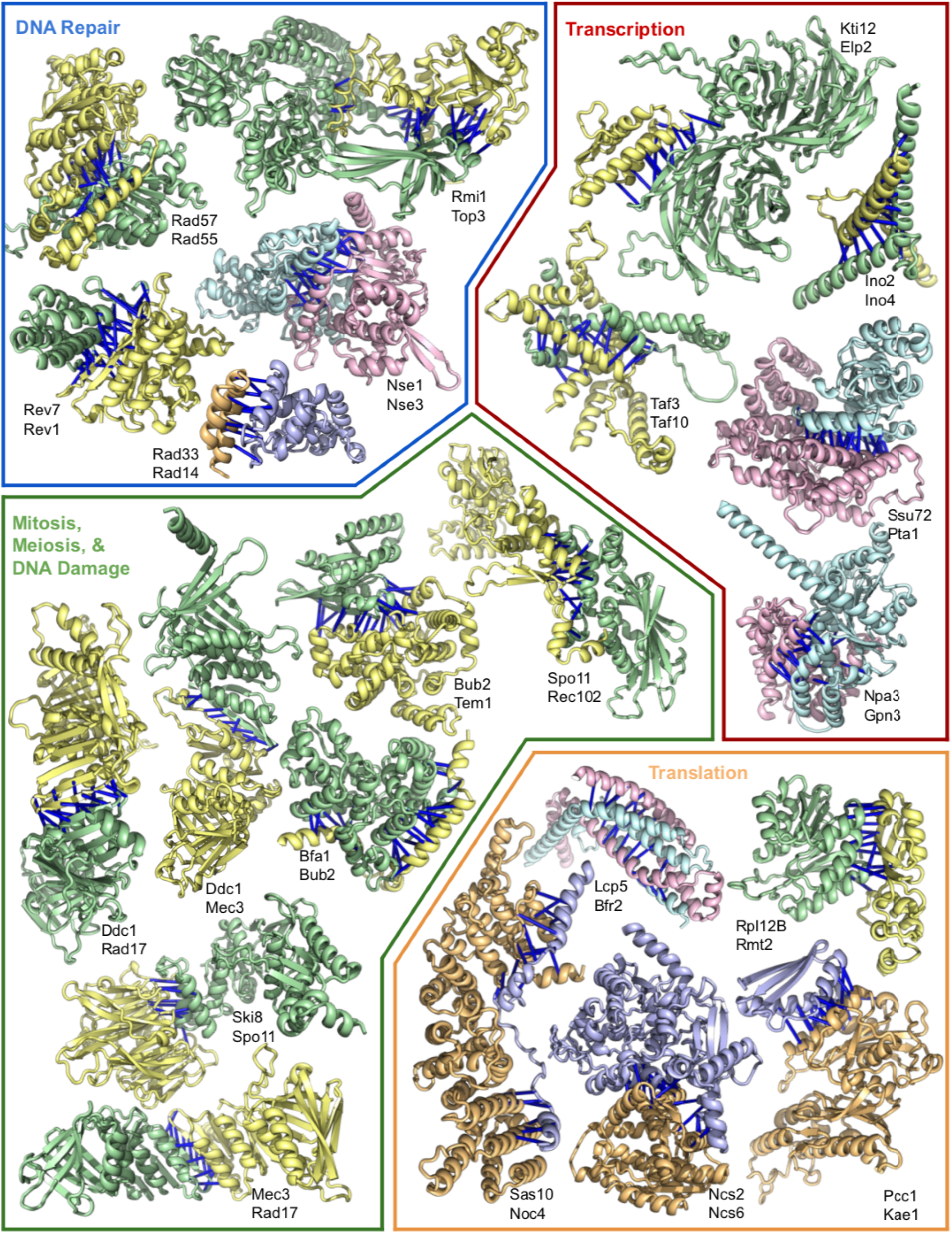
Protein complexes involved in transcription, translation, and DNA repair. Top predicted residue-residue contacts are indicated with bars. Pair color indicates the method of identification from Fig. 1D; experiment-guided pairs are yellow and green, “*de novo”* pairs are blue and light-orange, and pairs present in both datasets are teal and pink. Full names of each protein are in table S2.

**Figure 3.**
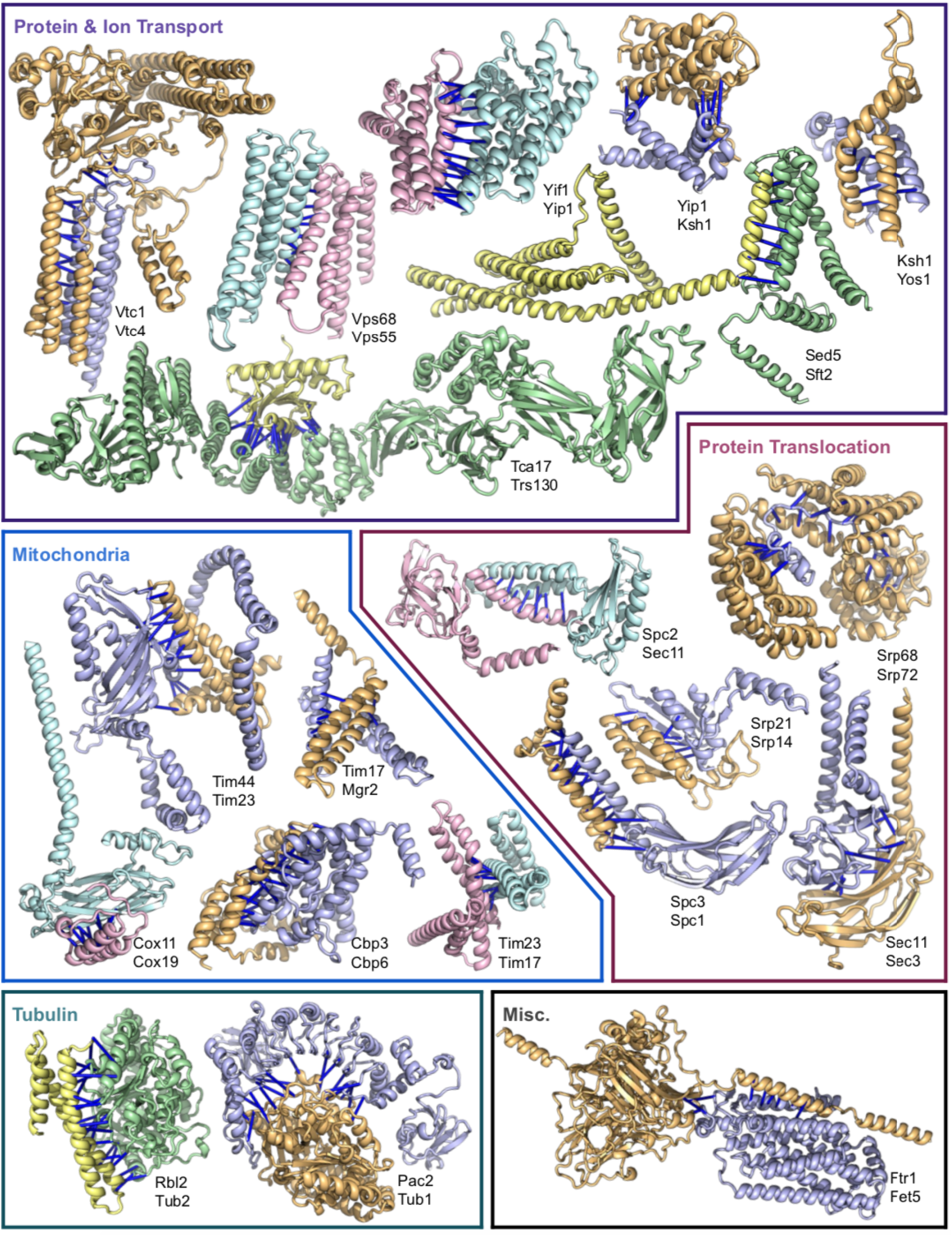
Protein complexes involved in protein transport, membrane translocation, and mitochondria. Bars and coloring as in Fig 2. Full names for each complex are in table S3.

**Figure 4.**
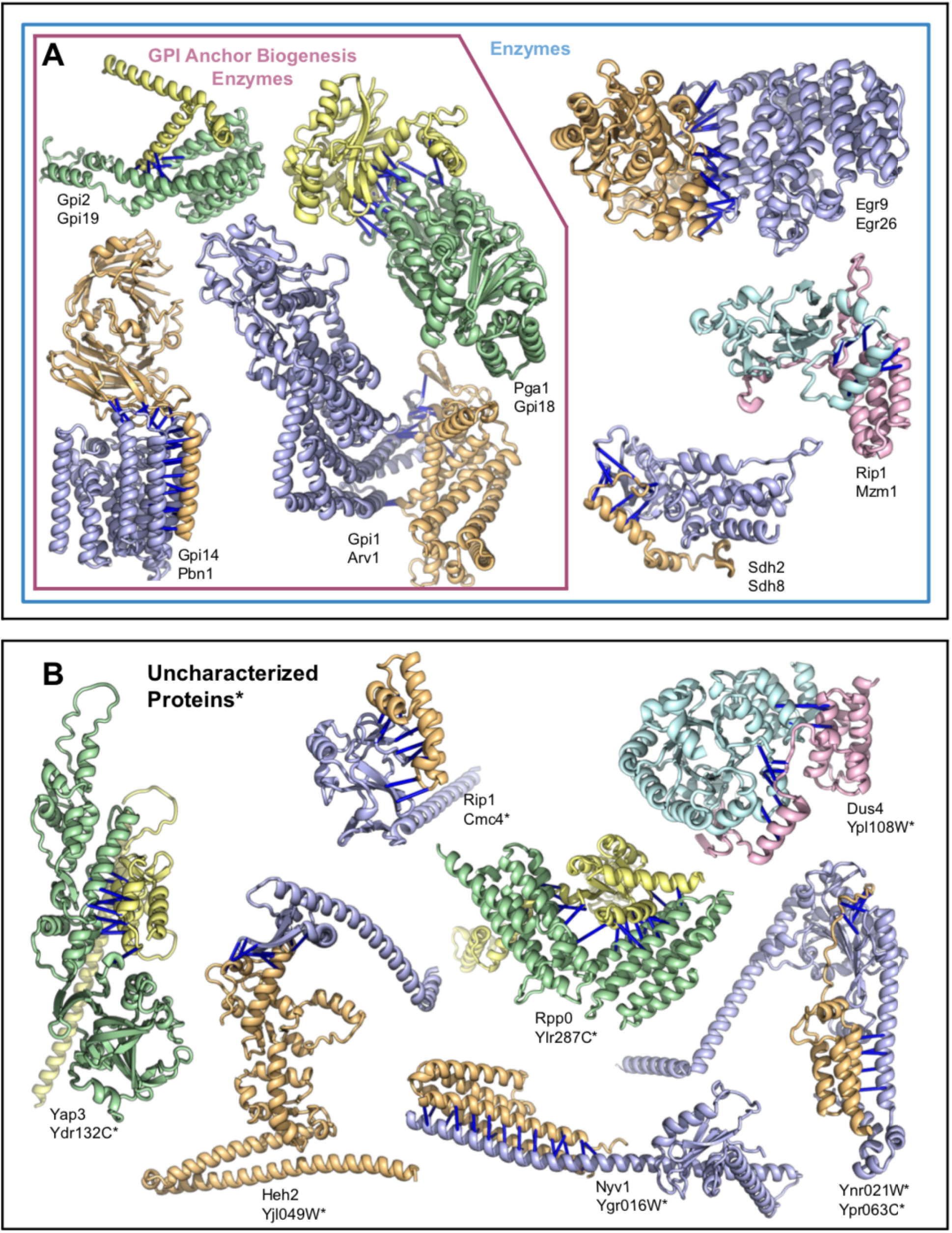
Protein complexes involved in metabolism, GPI anchor biosynthesis or including a protein of unknown function. Coloring is as in Fig. 2–3. Proteins annotated in the Uniprot database as uncharacterized proteins are denoted with an asterisk. Full names for each complex are in table S4.

**Figure 5.**
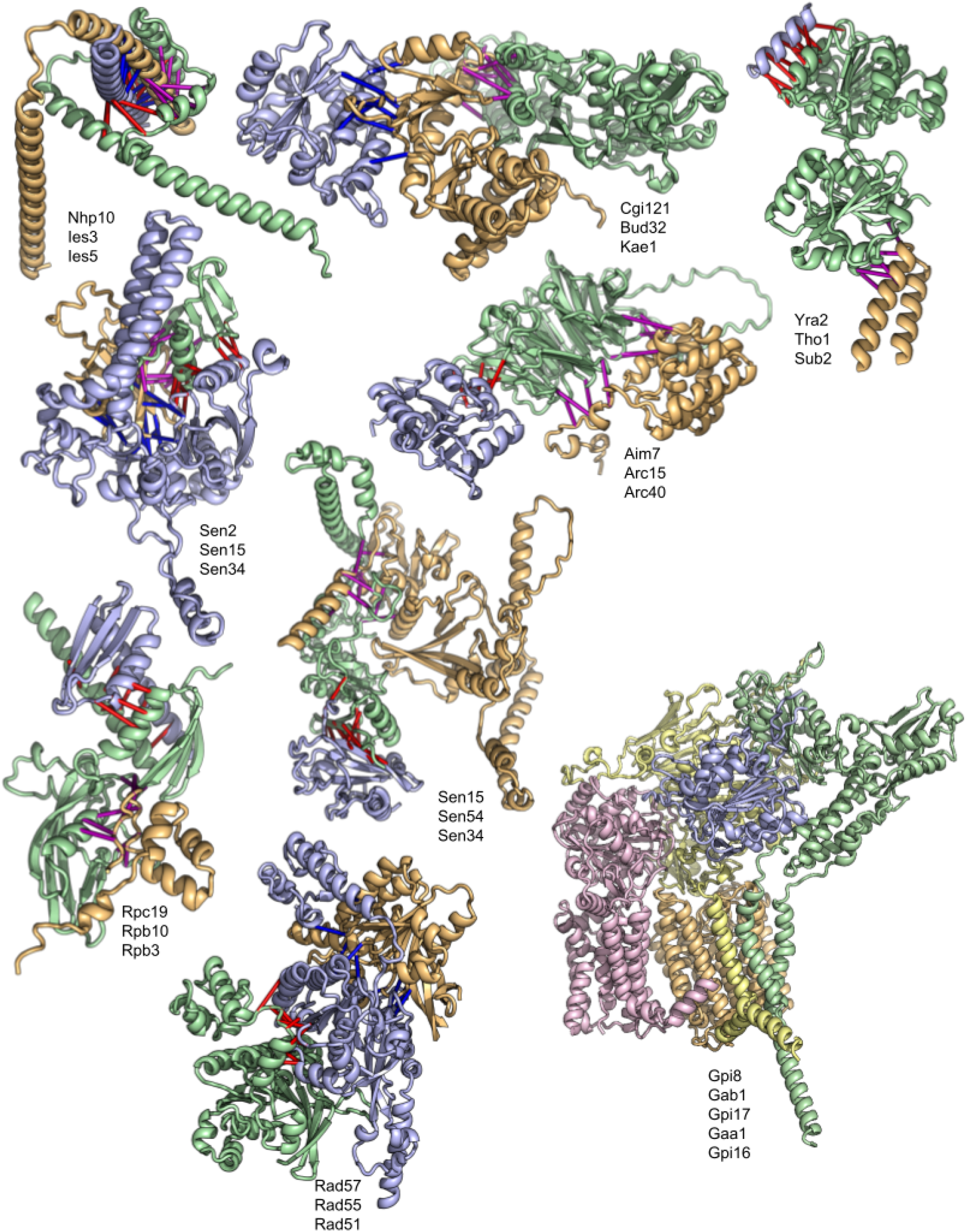
Higher order protein complexes. Top predicted residue-residue contacts for trimers are indicated with bars. Protein color indicates the protein number listed; protein 1 is blue, 2 is orange, 3 is green, 4 is pink, and 5 is light yellow. Bar color corresponds to the interacting protein pair; protein 1:2 are blue, 1:3 are red, 2:3 are purple. Full names of each protein within the complex are in table S5.

## Complexes involved in DNA homologous recombination and repair

The homologous recombination required for accurate chromosome segregation during meiosis is initiated by DNA double-strand breaks made by the Spo11 protein (*21*). Spo11 is essential for sexual reproduction in most Eukaryotes (*22*)(*23*), but mechanistic insight has been limited because of a deficit of high-resolution structural information. We predict the structures of complexes composed of Spo11 with its essential partners Ski8 and Rec102 (Fig. 2). The predicted Spo11–Ski8 complex structure is supported by crosslinking data and by the disruption of Spo11–Ski8 interactions *in vitro* and *in vivo* by mutations in Spo11 in the predicted interface regions (*24*)(*25*). Our model resembles a previous model based on the Ski3–Ski8 complex, with Ski8 contacting a sequence in Ski3 that is similar to the sequence QREIF_380_ in Spo11 (*25*)(*26*) (fig. S8A), but suggests a more extensive interaction surface than previously appreciated, involving an insertion in Ski8 that is present in *Saccharomyces* species but not in *Schizosaccharomyces pombe* and *Sordaria macrospora*, where Ski8 is also required for meiosis (*27*)(*28*) (fig. S8B,C). Rec102 was proposed from remote homology analysis to be the equivalent of the transducer domain of the Top6B subunit of archaeal topoisomerase VI (*29*), which couples ATP-dependent dimerization of a pair of Top6B subunits to DNA cleavage by a pair of Top6A subunits (*30*). Our predicted Rec102–Spo11 complex is similar to the Top6A–Top6B interface: a four-helix bundle consisting of two C-terminal helices from Rec102 and two helices from Spo11 (the first helix of the WHD plus a more N-terminally located helix) (fig. S8D). Alanine substitutions in this portion of Rec102 disrupt interaction with Spo11 and block meiotic recombination *in vivo* (*25*). The model clarifies the Spo11 portion of this interface, which was not well structured in previous homology models (*25*)(*29*). Both Rec102 and Top6B have long, helical arms that feed into the Spo11 interface; our model predicts a different angle for this arm and contains a previously unexpected kink that corresponds to a conserved sequence motif EYPMVF_192_ in *Saccharomyces* that is missing in both archaeal TopoVI and mammals (fig. S8D,E). Mutations in this region can suppress *rec104* conditional alleles (*31*), suggesting that this part of Rec102 is important for integrating Rec104 function into the Spo11 core complex.

Rad51 is a highly conserved DNA repair protein that carries out key reactions during homologous recombination; Rad51 paralogs are positive regulators of Rad51 activity (*32*), and mutations human paralogs are associated with Fanconi Anemia and multiple types of cancer (*33*). The yeast Rad51 paralogs Rad55 and Rad57 form a stable homodimer and the resulting Rad55–Rad57 complex accelerates assembly of Rad51 filaments on single–stranded DNA (ssDNA) during homologous recombination via a transient interaction with Rad51 (*34*). The lack of structural data for the Rad55–Rad57 complex and its interface with Rad51 has limited further mechanistic understanding of this process. We generated a model of the trimeric Rad55–Rad57-Rad51 complex, which in combination with the known Rad51 filament structure (*35*), suggests that Rad55–Rad57 binds at the 5’ end of the Rad51 filament where it could promote growth of the Rad51 filament in a directional manner (fig. S9).

## Complexes involved in translation and ribosome regulation

Throughout evolution the eukaryotic machinery for protein production has experienced a significant expansion in size and complexity (*36*), which facilitated the development of sophisticated mechanisms for the regulation of gene expression at the post-transcriptional level (*37*) and increased integration with the cellular environment (*38*). The expanded complexity of the eukaryotic translational machinery came at the cost of a highly complex process for ribosome maturation (*39*)). We generate models of complexes which had not been structurally characterized previously that involve components of translation apparatus (fig S10). Two complexes, Rpl12B–Rmt2 and Rpl7A–Fpr4, involve enzymes that introduce protein modifications such as arginine methylations or proline isomerizations (*40*) and suggest mechanisms to expand the chemical diversity of ribosomal proteins at functional sites (*41*)) which can play a role in regulating pathways in eukaryotes (*42*). A complex between components of the U3 ribosome-maturation factor and proteins involved in the regulation of glycerol, Lcp5–Sgd1 (*43*), could play a role in coupling translation with metabolism. A complex between eIF2B, an auxiliary factor for eIF2 recycling after GTP hydrolysis, and transcription factors of the STE12 family via DIG (DIG1/2) adaptor proteins could help couple translation and transcription: the delivery of the first aminoacyl-tRNA (Met-tRNA_i_^Met^) is a key event in eukaryotic translation regulated by the GTPase eIF2 (*44*) and targeting eIF2 via its nucleotide exchanger eIF2B is a basal mechanism of translation regulation that this complex suggests also is impacted by transcription factors. This possible cross-talk between ribosome-maturation pathways and metabolic sensors and translation initiation regulators such as eIF2 with transcription factors suggests exciting new avenues to further map the highly integrated nature of translation within eukaryotic cells.

## Complexes involving SUMO and ubiquitin ligases

Reversible covalent modifications of proteins with ubiquitin and the small ubiquitin-like modifier (SUMO) modulate protein-protein interactions, cellular localization, and stability (*45*). SUMO E3 ligases facilitate SUMO transfer, and Siz1, Siz2, Mms21, and Zip3 are the known SUMO ligases in budding yeast (*45*). We predict the structure of the Siz2 and Mms21 SUMO ligase complex (fig. S11A). Based on this model, it suggests that both E3s could act jointly to modify DNA associated substrates perhaps through the DNA binding SAP domain of Siz2 (*46*) or involving the Mms21 (alternative name: Nse2) containing Smc5–6 complex which modulates DNA recombination, replication and repair (*47*)(*48*). The Smc5–6 complex contains another RING-finger E3 ligase-like subunit, Nse1 (*49*) that interacts with Nse3 and Nse4. Our model of the yeast Nse1–Nse3–Nse4 complex (fig. S11B) is similar to a structure determined for the *Xenopus laevis* complex, despite the sequences of the yeast and Xenopus proteins being too distant for similarity to be detectable by BLAST.

SUMO-targeted ubiquitin ligases (STUbLs) are ubiquitin ligases that recognize SUMO-modified proteins. A STUbL consisting of the Slx8 ubiquitin ligase and the associated protein Slx5 functions in proteasome-mediated turnover of several proteins associated with DNA replication, repair and chromosome structure (*50*)(*51*)(*52*), but mechanistic details are currently lacking. We generated a lower confidence but intriguing model of a previously undescribed complex between Slx8 and Cue3 (Coupling of ubiquitin conjugation to endoplasmic reticulum (ER) degradation protein 3) (fig. S11C), possibly linking ubiquitination of substrates to degradation in ER.

## Complexes involved in chromosome segregation

The heterodecameric complex DASH/Dam1 (Dam1c) is composed of 10 proteins: Ask1, Dad1, Dad2, Dad3, Dad4, Dam1, Duo1, Hsk3, Spc19, and Spc34 which come together to form a “T” shape, and can further oligomerize into rings (*53*)((*54*)). During mitosis, these heterodecamers strengthen the attachment between kinetochores and microtubules, helping to ensure that chromosomes are accurately and faithfully segregated during cell division (*55*) by oligomerizing to form either partial or complete rings around microtubules and further contacting kinetochore components ((*56*))(*57*)(*58*). Microtubules are required for in-vivo ring formation, but a structure of the Dam1c ring complex from *Chaetomium themophilum* was determined in the absence of microtubules using monovalent salts (*59*). We generated structure models of nine heterodimeric complexes that encompass several members of Dam1c (fig. S12) that are largely consistent with the Dam1c structure, suggesting that the findings from the thermophile structure can likely be extended to *S. cerevisiae.* We go beyond the previous structure by predicting the structure of a potential inter-decamer interaction between Spc19 and Dad1 involving a flexible loop of Spc19 and the N-terminal region of Dad1 which could indicate how ring formation may take place in-vivo. Interestingly, these regions have previously been proposed as an interface between Dam1c decamers when forming a ring (*59*)). Our model predicts several electrostatic interactions (Spc19 LYS44 & Dad1 THR16; Spc19 LYS44 & Dad1 THR15; Spc19 ARG48 & Dad1 GLN23; Spc19 LYS44 & Dad1 TYR19) and various hydrophobic interactions (Spc19 VAL46 & Dad1 PHE20; Spc19 VAL46 & Dad1 TYR19) with high confidence, as well as a number of others which could drive this interaction.

## Complexes involved in membrane trafficking and protein transport

The ESCRT-III complex is involved in a number of cellular membrane remodeling pathways, including receptor downregulation, membrane repair, and cell division. Our predicted structure of the interface between the Vps2 and Vps24 subunits of the ESCRT-III complex resembles the polymerization interface of a different ESCRT-III subunit Snf7 (*60*), providing insight into the roles of these previously uncharacterized ESCRT-III subunits, and highlighting the generality of this mode of interaction in ESCRT-III complexes. Notably, previously unpublished mutations in Vps24 that prevent ESCRT function in multivesicular body sorting are located near the predicted interface between Vps2 and Vps24, supporting our model and the functional importance of the Vps2-Vps24 interaction (fig. S13). Vps55 and Vps68 are two conserved membrane proteins that are also important for endosomal cargo sorting; our predicted structure of their interaction provides possible clues about mechanism (*61*).

SNARE proteins drive intracellular membrane fusion reactions between transport vesicles and organelles. Our prediction of the structure of a complex between the SNARE Sed5 and the transmembrane protein of unknown function Sft2 unexpectedly predicted an interaction between transmembrane domains of the two proteins (fig. S14). SNARE localization is thought to occur via interactions of cytoplasmic domains with cytoplasmic sorting factors, but this prediction, together with genetic evidence (*62*), suggests SNARE localization or function may be subject to additional mechanisms via interactions with transmembrane protein regulators. The small membrane protein Ksh1 is conserved across eukaryotes, essential for growth, and plays an unknown role in secretion (*63*). We predicted structures of complexes between Ksh1 with Yos1 and Yip1, membrane proteins reported to form a complex that also includes Yif1 and interacts with Rab GTPases (*64*) (Fig 3). These structures suggest Ksh1 is a fourth member of this enigmatic complex essential to the secretory pathway, and explains how Ksh1 can play a role in secretion despite its small size of 72 amino acids.

The GARP complex is a multisubunit tethering complex (MTC) that mediates docking and fusion of vesicles with the Golgi apparatus (*65*). Our approach generated predictions that included each of the four GARP subunits, and we generated models for the entire complex (fig. S15A). In this model, the four subunits assemble through a four-helix bundle. In each of the three larger subunits, Vps52, Vps53, and Vps54, C-terminal domains comprising “CATCHR” folds emanate from the bundle. This architecture resembles portions of the cryo-EM structure of the Exocyst complex, a distinct MTC that mediates fusion of vesicles at the plasma membrane (*66*), which possesses two separate four-helix bundles organizing its eight subunits. In our prediction, the GARP subunit CATCHR domains appear to be somewhat flexibly linked to the central four-helix bundle, and hence we overlaid the structure predictions for Vps52, Vps53, and Vps54 onto the central four-helix bundle (fig. S15B). The resulting model has a striking resemblance to previously published 2D classes (fig. S15C) from a negative-stain EM analysis of the GARP complex (*67*). These predictions will facilitate structure-guided experiments to elucidate the mechanism of MTC function.

The yeast TRAPPII complex is a conserved activator of Rab11, comprising 22 subunits encoded by 10 genes (*68*). Our prediction of the structure of the Tca17–Trs130 subunit interface (Fig. 3) is essentially identical to that obtained independently by preliminary cryo-EM analysis (unpublished data), further attesting to the accuracy of our complex predictions. The vacuolar transporter chaperone (VTC) is a 5-subunit complex that synthesizes polyphosphate to regulate cellular phosphate levels (*69*). Structures are only known for some soluble portions of this complex, including the catalytic domain of the Vtc4 subunit (*70*). Our model of the previously non-structurally characterized Vtc1–Vtc4 subcomplex suggests that the cytosolic active site is positioned by the complex to feed the polyphosphate product through a membrane pore into the lumen of the lysosome (fig. S16).

Golgi-resident protein, Grh1, forms a tethering complex with Uso1 and Bug1 that interacts with the COPII coat protein complex, Sec23/Sec24. The tether is thought to participate in COPII vesicle capture (*71*)(*72*), although the mechanism remains unclear. The C-terminus of Grh1 contains a predicted intrinsically disordered region (IDR) with a net positively charged cluster and a triple-proline motif (fig. 17A). Our model of the Sec23–Grh1 complex contains an interface between the Sec23 gelsolin domain and the PPP motif of Grh1 (*73*), and an interface between the Grh1 IDR and Sec23 involving a disorder-to-helical transition (fig. S17B). A similar multivalent interface also drives interaction between Sec23 and the COPII coat scaffolding protein, Sec31 (*74*). Our structure of the complex suggests that the combinatorial multivalent interaction between Grh1 and Sec23 may compete with the interaction between Sec31 and Sec23 to promote vesicle uncoating; consistent with this model, Grh1 is recruited to GST-Sec23, dependent on the IDR, and competes for Sec31 binding (fig. S17C).

## Complexes involved in glycolipid anchor attachment to proteins

Glycosylphosphatidylinositol transamidase (GPI-T) is a pentameric enzyme complex of unknown structure (*75*)(*76*) which catalyzes the attachment of GPI anchors to the C-terminus of specific substrate proteins, based on recognition of a nondescript C-terminal signal peptide (*77*). GPI-T catalyzes the removal of this signal sequence, replacing it with a new amide bond to an ethanolamine phosphate in the GPI anchor. The five subunits of *S. cerevisiae* GPI-T are Gpi8 (the catalytic active site), Gpi16, Gaa1, Gpi17, and Gab1 (*78*)(*79*)(*75*). Our large scale modeling approach generated models for the pairwise interactions between Gpi8 and Gpi17; Gab1 and Gaa1; Gab1 and Gpi17; and Gaa1 and Gpi16, and we subsequently assembled models of the full-length, pentameric GPI-T in one shot starting from the sequences of all components (Fig. 5). Several features of this model are consistent with previous characterization of this enzyme. *S. cerevisiae* GPI-T can be purified as a core heterotrimer, containing only Gpi8, Gpi16, and Gaa1 (*78*); our GPI-T model confirms extensive interactions between the soluble domains of these three subunits. This model also recapitulates the disulfide bond between Gpi8 (Cys85) and Gpi16 (Cys202), previously characterized for the human GPI-T (*80*) (the existence of this disulfide bond in the yeast GPI-T has been called into question (*81*).) Gaa1 is essential for binding of the GPI anchor to GPI-T (*82*) and the hydrophobic Gab1 is also predicted to participate in anchor recognition (*75*). Our model positions the transmembrane regions of Gaa1 and Gab1 against each other. The catalytic dyad in Gpi8 (Cys199 and His157) faces these transmembrane domains, and abuts directly against a highly conserved face of Gaa1, proposed to recognize the GPI anchor glycans (*83*)(*84*). In our model, these three subunits interact in a way consistent with binding of the GPI anchor to position the modifying amine in the Gpi8 active site for catalysis. Gpi16 is immediately adjacent to these interactions and is likely to also be involved in anchor recognition. In vivo, GPI-T is likely to be a dimer of pentamers, with dimerization occurring on one face of the caspase-like Gpi8 subunit (*78*)(*85*)(*84*). This decameric complex was too large for us to model computationally; however the pentameric complex we present here leaves open the dimerization face of Gpi8, consistent with probable dimerization. It also suggests that Gaa1 and Gpi17 would also participate in dimerization of this enzyme. The functional role of Gpi17 has been elusive, but our model now suggests Gpi17 collaborates with Gpi8 and Gpi16 to form a channel through which the C-terminal GPI-T signal peptide might be threaded (fig. S18). In humans, mutations in GPI-T subunits are associated with and also cause neurodevelopmental disorders (*86*). Each subunit contributes to different cancer mechanisms, in some cases by perturbing GPI anchoring of specific receptors and in others by separating from GPI-T to alter disparate signal transduction pathways (*76*). Now, with a structural model in hand, these mechanisms can be examined at a molecular level.

## Conclusion

Our approach extends the range of large scale deep learning based structure modeling from monomeric proteins to protein assemblies. As highlighted by the above examples, following up on the many new complexes presented here should advance understanding of a wide range of eukaryotic cellular processes and provide new targets for therapeutic intervention. The methods can be extended directly to large scale mapping of interactions in the human proteome, but considerably more compute time will be required given the much larger number of possible pairwise interactions, coevolutionary signal will be weaker for the subset of human proteins that are unique to higher eukaryotes since there are smaller numbers of homologous sequences, and it will be more difficult to distinguish interacting partners for closely related paralogs due to more extensive gene duplication. However, investigating interactions of individual proteins or subsets of proteins, for example, deorphanization of orphan receptors, should be immediately accessible using our approach. Training RF and AF on protein complexes should further improve performance of both methods, particularly for protein pairs with shallower pMSAs or weaker and more transient interaction, and could reduce the dependence on ortholog identification. Together with the advances in monomeric structure prediction, our results herald a new era of structural biology in which computation plays a fundamental role in both interaction discovery and structure determination.

## Acknowledgements

We thank Eric Horvitz, Nick V. Grishin, Hahnboem Park, and James H. Thomas for helpful discussions, and Luki Goldschmidt and Aaron Guillory for computing resource management. Additionally, we are grateful to Martin Bard, Trisha N Davis, David G Drubin, Maitreya J Dunham, Scott Emr, Frederick Hughson, James Hurley, Kenji Murakami, Nobuhiro Nakamura, Eva Nogales, Randy Schekman, Shu-ou Shan, Soyeon Showman, Kaoru Sugasawa, and Sho Suzuki for their correspondence and biological expertise.

## Author contributions

QC and DB conceived the research; JP and QC prepared the sequence alignments used in the screen; MB and SO implemented the RoseTTAFold pipeline and repurposed AlphaFold for complex modeling; JP, JZ, and QC designed the computational pipeline; IRH, MB, IA, and QC carried out the screen; IRH, AK, and QC analyzed and presented the results; IRH, AK, QC and DB coordinated the collaborative efforts; TJN, SB, SB, VGS, XHL, KL, ZZ, DB, UR, ISF, BS, DJB, ECG, SB, SK, EAM, JCF, and TLM provided biological insights on specific examples; IRH, MB, AK, QC, and DB wrote the manuscript; all authors discussed the results and commented on the manuscript.

## Supplemental Figures

**Figure S1.**
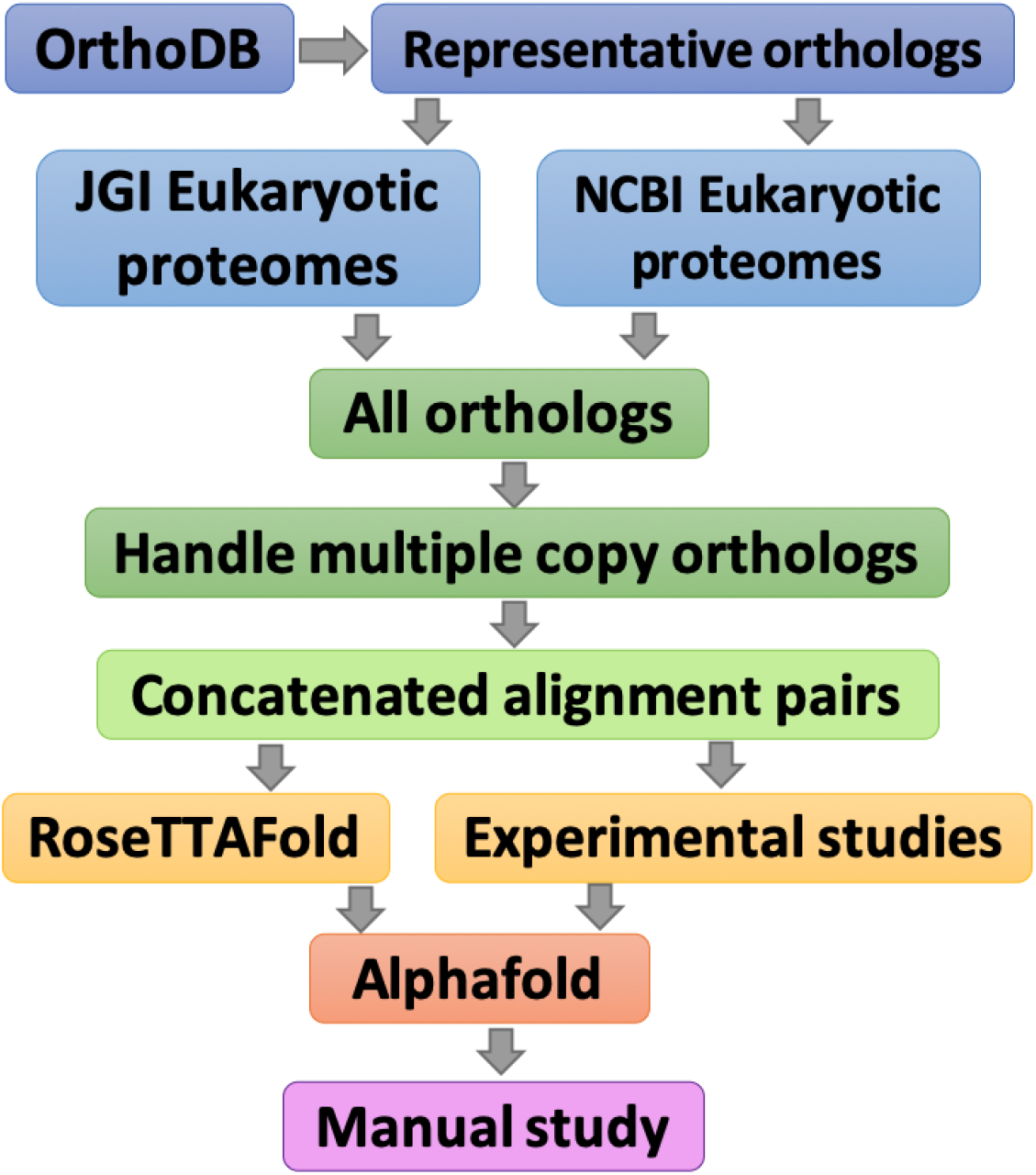
Flowchart of the *in silico* PPI screen pipeline.

**Figure S2.**
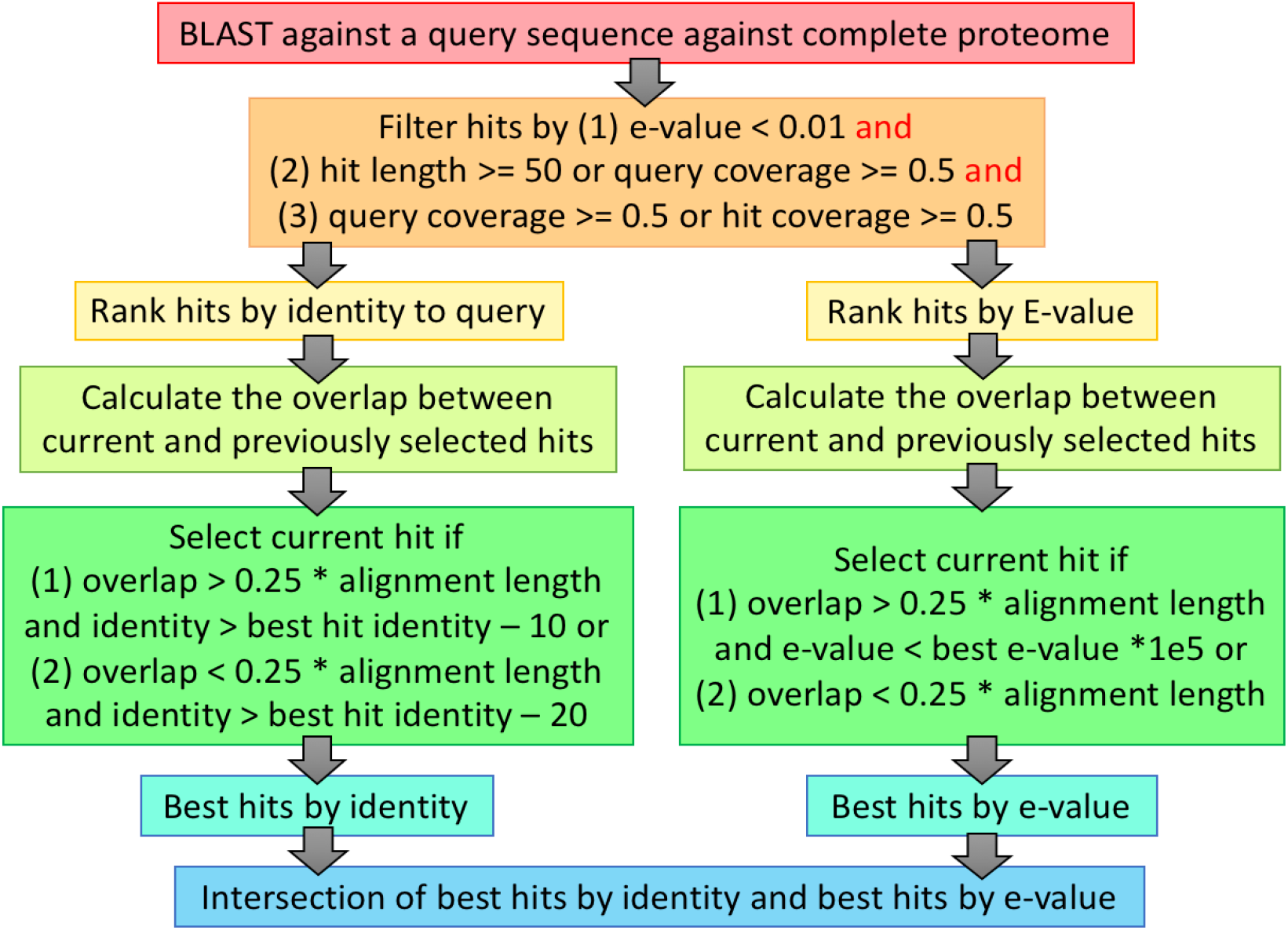
Procedure for identifying best hit to a query protein in the complete proteome of a different species.

**Figure S3.**
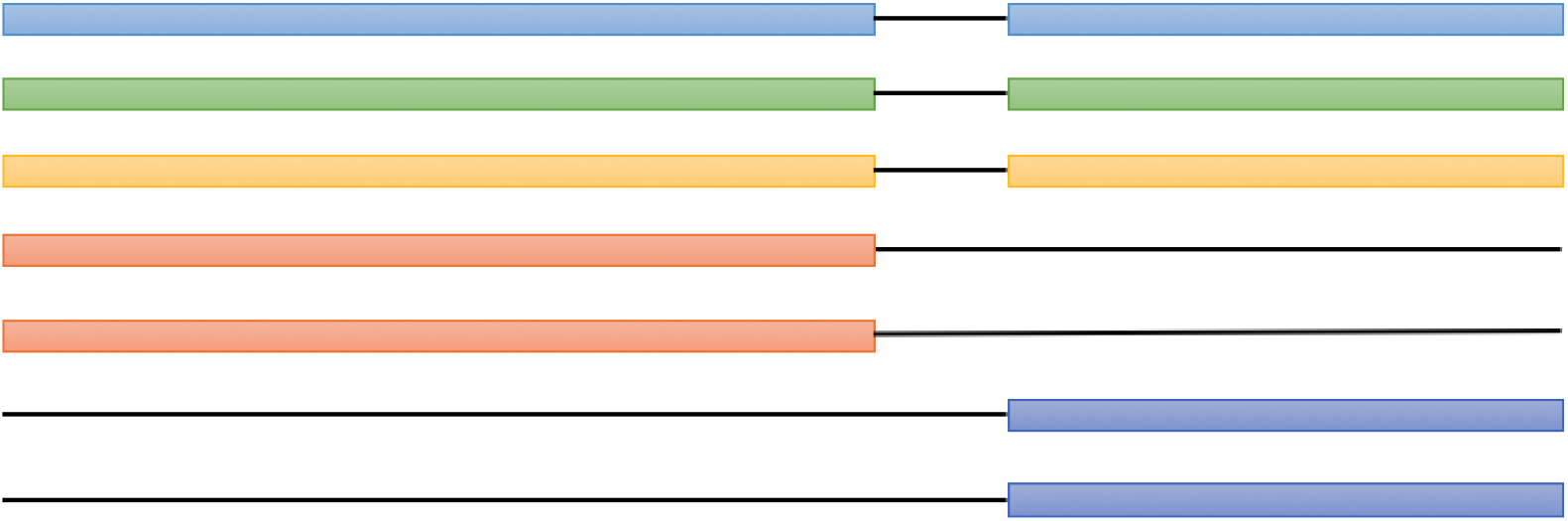
Protocol for generating pMSA. Proteins from the same proteome (the same organism) are colored with the same color, and a black line indicates gaps. We did not explicitly add gaps between two proteins, but we introduced a gap in the residue numbers at the junction between the two proteins when we analyzed the pMSA by RoseTTAFold or AlphaFold.

**Figure S4.**
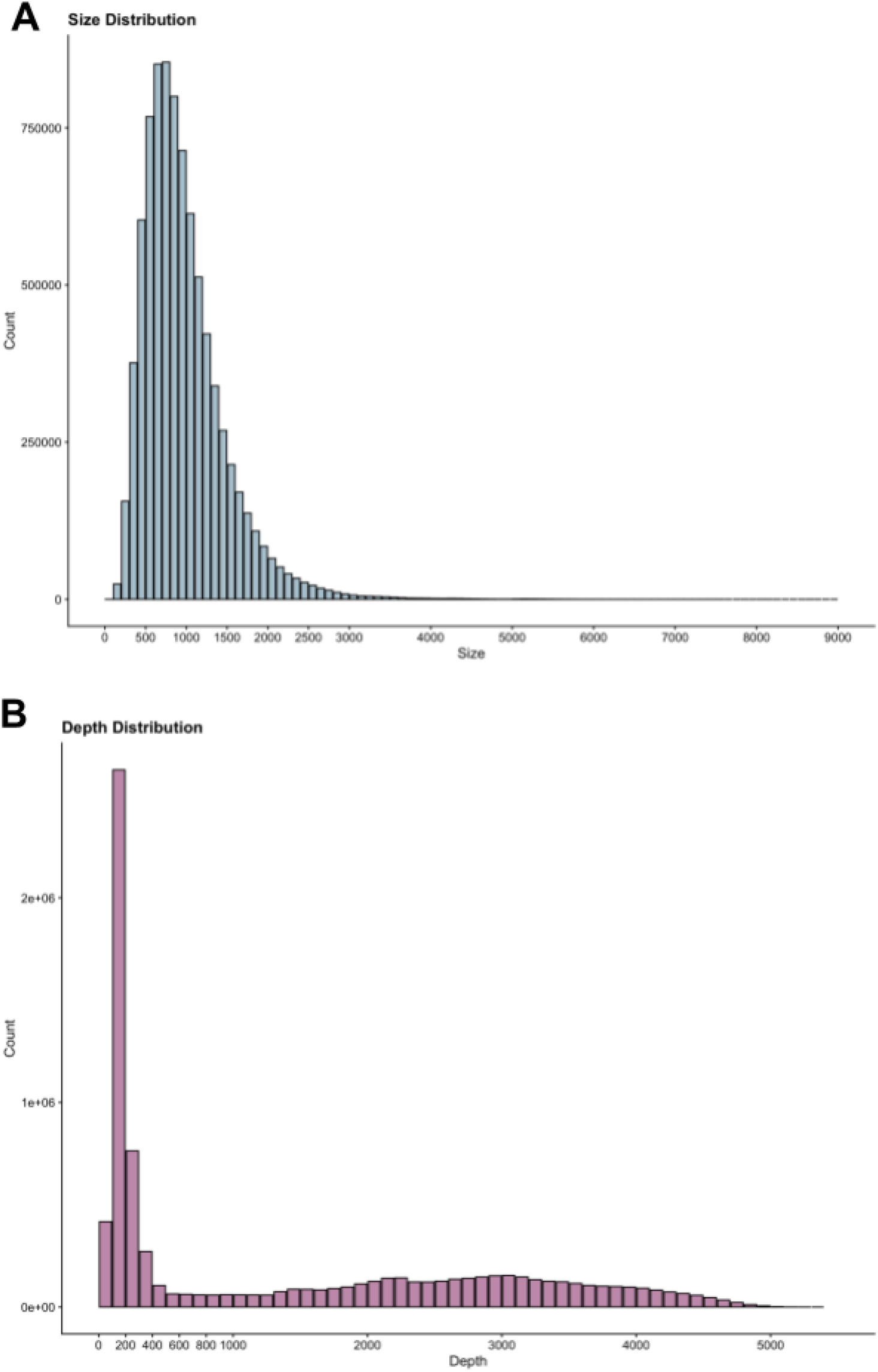
Length (A) and depth (B) distributions of pMSA.

**Figure S5.**
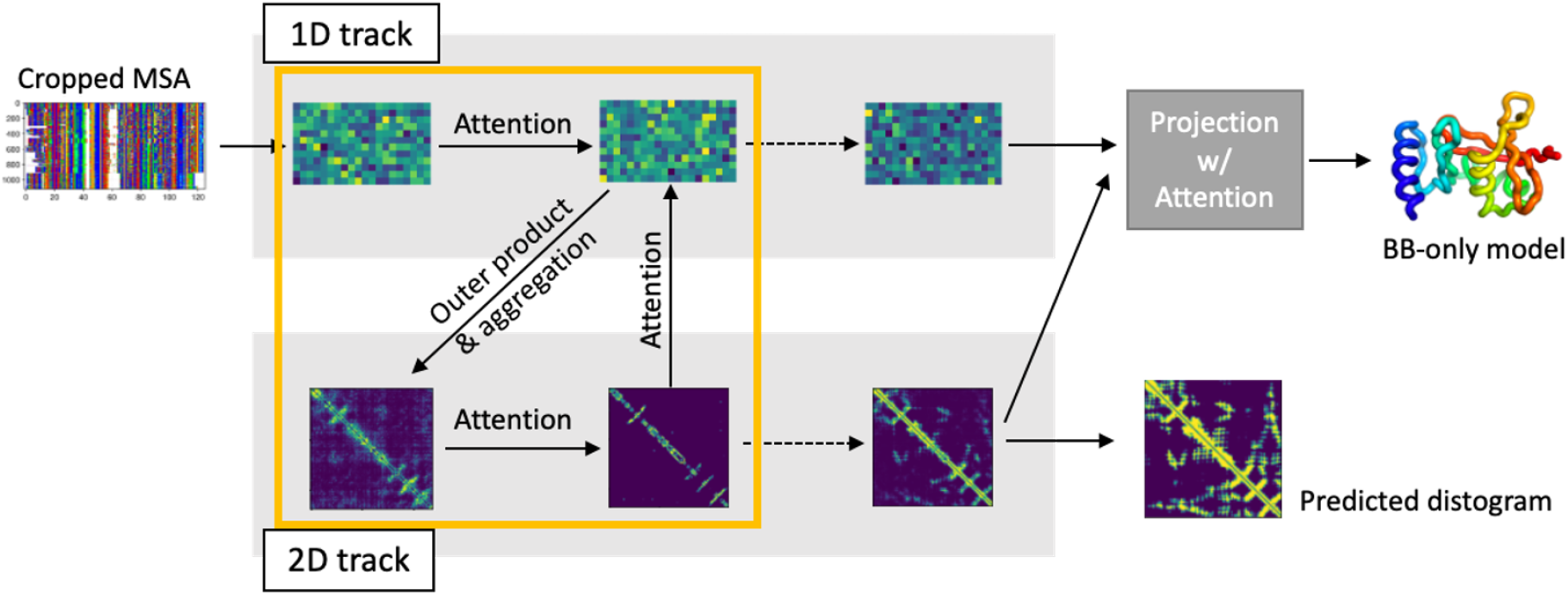
A lighter-weight RoseTTAFold two-track (RF2t) model.

**Table S1.** Datasets we obtained from Yeast Interactome Database.

| Dataset | Description |
| --- | --- |
| Y2H-union | The union of CCSB-YI1, Ito-core, and Uetz-screen |
| Combined-AP/MS | Co-complex membership associations |
| LC-multiple | Literature-curated interactions |
| PRS | Positive Reference Set |
| Binary-GS | Binary Gold standard Set |

**Table S2.** Extended annotations for modeled PPIs in Fig. 2.

| **Protein 1** | **Protein 2** | **Group** | **Protein 1 Annotation** | **Protein 2 Annotation** |
| --- | --- | --- | --- | --- |
| Nse1 | Nse3 | DNA Repair | Non-structural maintenance of chromosomes element 1 | Non-structural maintenance of chromosome element 3 |
| Rad33 | Rad14 | DNA Repair | DNA repair protein RAD33 | DNA repair protein RAD14 |
| Rmi1 | Top3 | DNA Repair | RecQ-mediated genome instability protein 1 | DNA topoisomerase 3 |
| Rev7 | Rev1 | DNA Repair | DNA polymerase zeta processivity subunit | DNA repair protein REV1 |
| Rad57 | Rad55 | DNA Repair | DNA repair protein RAD57 | DNA repair protein RAD55 |
| Ddc1 | Rad17 | Mitosis, Meiosis, or DNA Damage | DNA damage checkpoint protein 1 | DNA damage checkpoint control protein RAD17 |
| Ddc1 | Mec3 | Mitosis, Meiosis, or DNA Damage | DNA damage checkpoint protein 1 | DNA damage checkpoint control protein MEC3 |
| Mec3 | Rad17 | Mitosis, Meiosis, or DNA Damage | DNA damage checkpoint control protein MEC3 | DNA damage checkpoint control protein RAD17 |
| Spo11 | Rec102 | Mitosis, Meiosis, or DNA Damage | Meiosis-specific protein SPO11 | Meiotic recombination protein REC102 |
| Ski8 | Spo11 | Mitosis, Meiosis, or DNA Damage | Antiviral protein SKI8 | Meiosis-specific protein SPO11 |
| Bub2 | Tem1 | Mitosis, Meiosis, or DNA Damage | Mitotic check point protein BUB2 | Protein TEM1 |
| Bfa1 | Bub2 | Mitosis, Meiosis, or DNA Damage | Mitotic check point protein BFA1 (Cell cycle arrest protein) | Mitotic check point protein BUB2 |
| Taf3 | Taf10 | Transcription | Transcription initiation factor TFIID subunit 3 | Transcription initiation factor TFIID subunit 10 |
| Ssu72 | Pta1 | Transcription | RNA polymerase II subunit A C-terminal domain phosphatase SSU72 | Pre-tRNA-processing protein PTA1 |
| Npa3 | Gpn3 | Transcription | GPN-loop GTPase 1 | GPN-loop GTPase 3 |
| Ino2 | Ino4 | Transcription | Protein INO2 | Protein INO4 |
| Kti12 | Elp2 | Transcription | Protein KTI12 (Gamma-toxin target protein 4) (Killer toxin insensitivity protein 12) | Elongator complex protein 2 (Gamma-toxin target 2) |
| Sas10 | Noc4 | Translation | Something about silencing protein 10 | Nucleolar complex protein 4 |
| Lcp5 | Bfr2 | Translation | U3 small nucleolar ribonucleoprotein protein LCP5 | Protein BFR2 (Brefeldin A resistance protein 2) |
| Pcc1 | Kae1 | Translation | EKC/KEOPS complex subunit PCC1 (Polarized growth chromatin-associated controller 1) | tRNA N6-adenosine threonylcarbamoyltransferase |
| Ncs2 | Ncs6 | Translation | Cytoplasmic tRNA 2-thiolation protein 2 | Cytoplasmic tRNA 2-thiolation protein 1 |
| Rpl12B | Rmt2 | Translation | 60S ribosomal protein L12-B | Protein arginine N-methyltransferase 2 |

**Table S3.** Extended annotations for modeled PPIs from Fig. 3.

| **Protein 1** | **Protein 2** | **Group** | **Protein 1 Annotation** | **Protein 2 Annotation** |
| --- | --- | --- | --- | --- |
| Yif1 | Yip1 | Protein & Ion Transport | Protein transport protein YIF1 | Protein transport protein YIP1 |
| Ksh1 | Yos1 | Protein & Ion Transport | Protein Kish | Protein transport protein YOS1 |
| Tca17 | Trs130 | Protein & Ion Transport | TRAPP-associated protein TCA17 | Trafficking protein particle complex II-specific subunit 130 |
| Vtc1 | Vtc4 | Protein & Ion Transport | Vacuolar transport chaperone 1 | Vacuolar transporter chaperone 4 |
| Vps68 | Vps55 | Protein & Ion Transport | Vacuolar protein sorting-associated protein 68 | Vacuolar protein sorting-associated protein 55 |
| Sed5 | Sft2 | Protein & Ion Transport | Integral membrane protein SED5 | Protein transport protein SFT2 |
| Yip1 | Ksh1 | Protein & Ion Transport | Protein transport protein YIP1 | Protein kish |
| Cbp3 | Cbp6 | Mitochondria | Protein CBP3, mitochondrial | Cytochrome B pre-mRNA-processing protein 6 |
| Cox11 | Cox19 | Mitochondria | Cytochrome c oxidase assembly protein COX11, mitochondrial | Cytochrome c oxidase assembly protein COX19 |
| Tim17 | Mgr2 | Mitochondria | Mitochondrial import inner membrane translocase subunit TIM17 | Protein MGR2 (Mitochondrial genome-required protein 2) |
| Tim44 | Tim23 | Mitochondria | Mitochondrial import inner membrane translocase subunit TIM44 | Mitochondrial import inner membrane translocase subunit TIM23 |
| Tim23 | Tim17 | Mitochondria | Mitochondrial import inner membrane translocase subunit TIM23 | Mitochondrial import inner membrane translocase subunit TIM17 |
| Spc3 | Spc1 | Protein Translocation | Signal peptidase complex subunit SPC3 | Signal peptidase complex subunit SPC1 |
| Sec11 | Spc3 | Protein Translocation | Signal peptidase complex catalytic subunit SEC11 | Signal peptidase complex subunit SPC3 |
| Spc2 | Sec11 | Protein Translocation | Signal peptidase complex subunit SPC2 | Signal peptidase complex catalytic subunit SEC11 |
| Srp68 | Srp72 | Protein Translocation | Signal recognition particle subunit SRP68 | Signal recognition particle subunit SRP72 |
| Srp21 | Srp14 | Protein Translocation | Signal recognition particle subunit SRP21 | Signal recognition particle subunit SRP14 |
| Pac2 | Tub1 | Protein Translocation | Protein PAC2 | Tubulin alpha-1 chain |
| Rbl2 | Tub2 | Protein Translocation | Tubulin-specific chaperone A | Tubulin beta chain |
| Ftr1 | Fet5 | Misc. | Plasma membrane iron permease | Iron transport multicopper oxidase FET5 |

**Table S4.** Extended annotations for modeled PPIs in Fig. 4.

| Protein 1 | Protein 2 | Group | Protein 1 Annotation | Protein 2 Annotation |
| --- | --- | --- | --- | --- |
| Erg9 | Erg26 | Enzyme | Squalene synthase | Sterol-4-alpha-carboxylate 3-dehydrogenase, decarboxylating |
| Rip1 | Mzm1 | Enzyme | Cytochrome b-c1 complex subunit Rieske, mitochondrial | Mitochondrial zinc maintenance protein 1, mitochondrial |
| Sdh2 | Sdh8 | Enzyme | Succinate dehydrogenase [ubiquinone] iron-sulfur subunit, mitochondrial | Succinate dehydrogenase assembly factor 4, mitochondrial |
| Gpi2 | Gpi9 | Enzyme (GPI) | Phosphatidylinositol N-acetylglucosaminyltransferase GPI2 subunit | Phosphatidylinositol N-acetylglucosaminyltransferase subunit GPI19 |
| Gpi14 | Pbn1 | Enzyme (GPI) | GPI mannosyltransferase 1 | Protein PBN1 (Protease B non-derepressible protein 1) |
| Gpi1 | Arv1 | Enzyme (GPI) | Phosphatidylinositol N-acetylglucosaminyltransferase subunit GPI1 | Protein ARV1 |
| Pga1 | Gpi18 | Enzyme (GPI) | GPI mannosyltransferase 2 subunit PGA1 | GPI mannosyltransferase 2 |
| Heh2 | Yjl049W | Uncharacterized | Inner nuclear membrane protein HEH2 | Uncharacterized protein YJL049W |
| Rip1 | Cmc4 | Uncharacterized | Cytochrome b-c1 complex subunit Rieske, mitochondrial | Cx9C motif-containing protein 4, mitochondrial |
| Rpp0 | Ylr287C | Uncharacterized | 60S acidic ribosomal protein P0 | Uncharacterized protein YLR287C |
| Dus4 | Ypl108W | Uncharacterized | tRNA-dihydrouridine(20a/20b) synthase [NAD(P)+] | Uncharacterized protein YPL108W |
| Ynr021W | Ypr063C | Uncharacterized | UPF0674 endoplasmic reticulum membrane protein YNR021W | Uncharacterized protein YPR063C |
| Nyv1 | Ygr016W | Uncharacterized | Vacuolar v-SNARE NYV1 | Uncharacterized membrane protein YGR016W |
| Yap3 | Ydr132C | Uncharacterized | AP-1-like transcription factor YAP3 | Uncharacterized protein YDR132C |

**Table S5.**
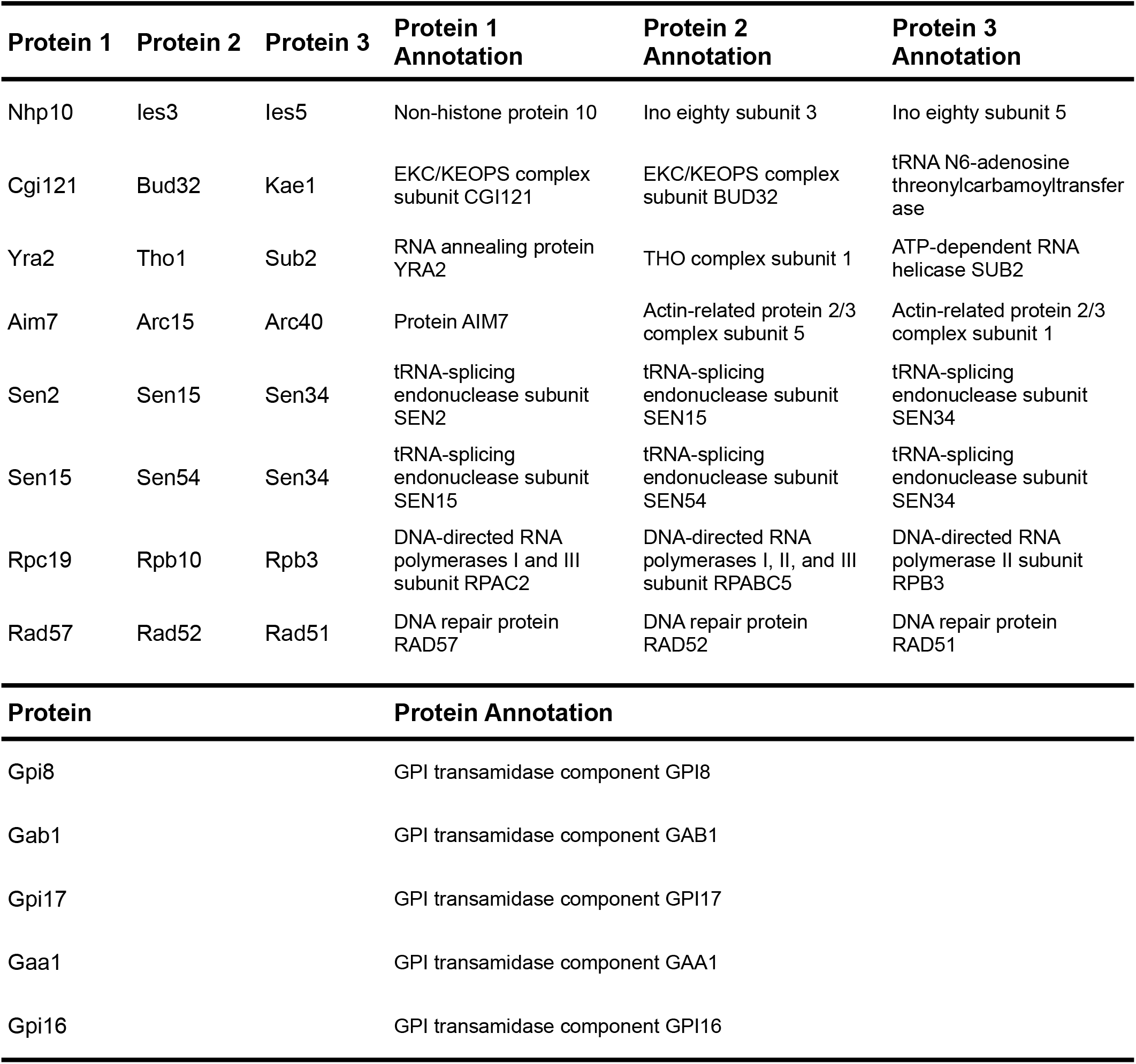
Extended annotations for modeled PPIs in Fig. 5.

**Figure S7.**
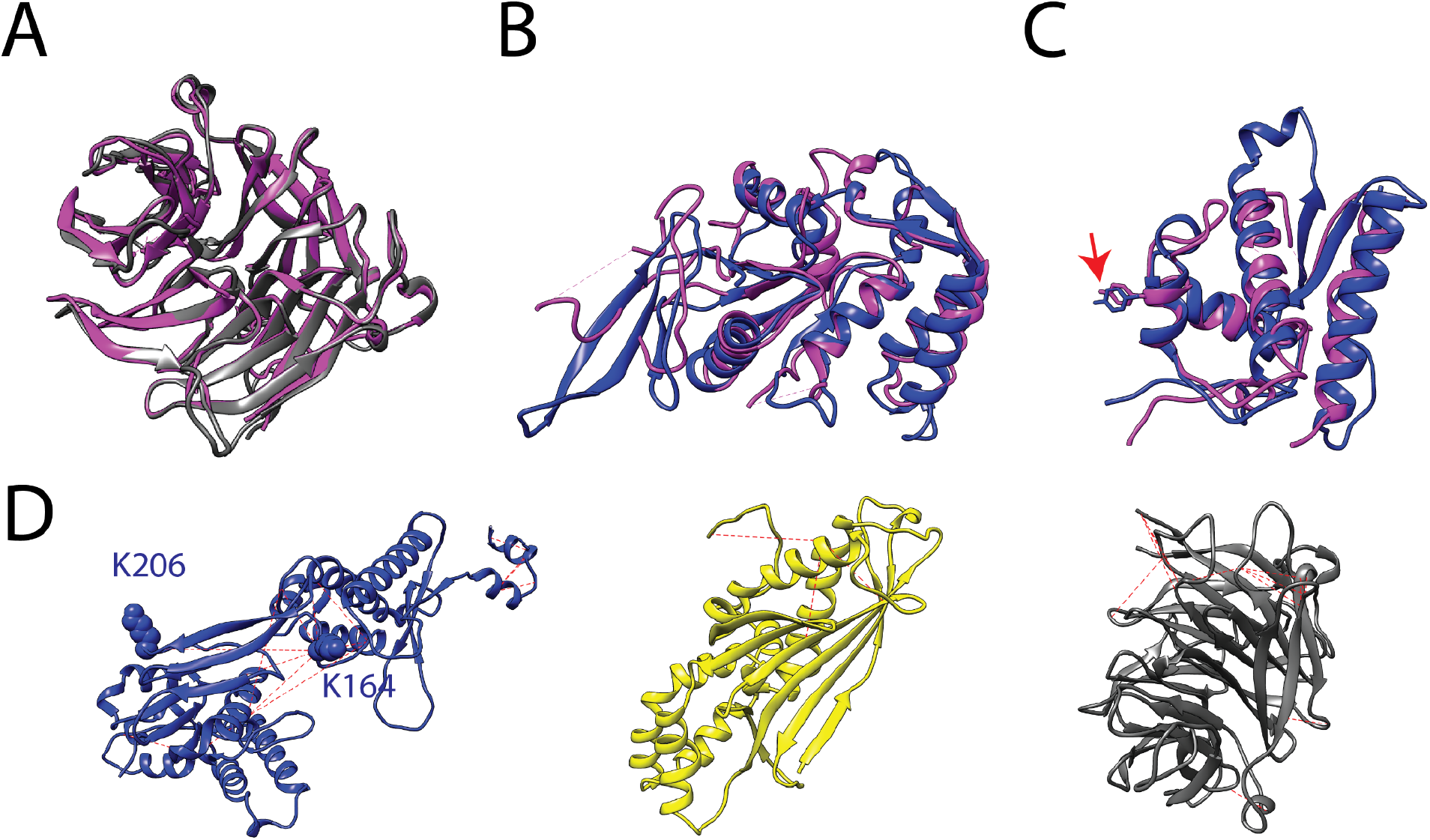
Ski8 complex monomer models. (**A**) Comparison of the AF model for Ski8 (gray) and a crystal structure (magenta, pdb: 1sq9 (*87*)). The root mean square deviation (RMSD) between the aligned structures is 1.1 Å. (**B**) Comparison of the predicted Spo11 Toprim domain structure (blue; residues 173 to 398) with the Toprim domain from *M. mazei* Top6A (magenta; residues 147 to 367 from pdb 2q2e (*88*)). The RMSD is 1.1 Å. (**C**) Comparison of the predicted WHD for Spo11 (blue; residues 24 to 172) with the WHD for *M. mazei* Top6A (magenta; residues 15 to 146). The RMSD is 1.2 Å. The catalytic tyrosines are positioned nearly identically for the two proteins (arrow). (**D**) Comparison of models to intramolecular crosslinking data for Spo11 (blue), Rec102 (yellow), and Ski8 (gray). Red dashed lines connect the acarbons of lysine pairs that were observed to be crosslinked in a recent study of the Spo11-Ski8-Rec102-Rec104 complex (*25*)). In the Spo11 model, 14 out of 15 cross-linked lysine pairs with high mass spectrometry counts (≥10) are within the crosslinker range limit of 27.4 Å. For the only exception (K164-K206, predicted distance of 30.7 Å), both lysines are in predicted loop regions which may be conformationally flexible. Similarly, all of the high-frequency crosslinked lysine pairs are within the crosslinking distance limit in the AF models for Rec102 (3 lysine pairs) and Ski8 (14 pairs).

**Figure S8.**
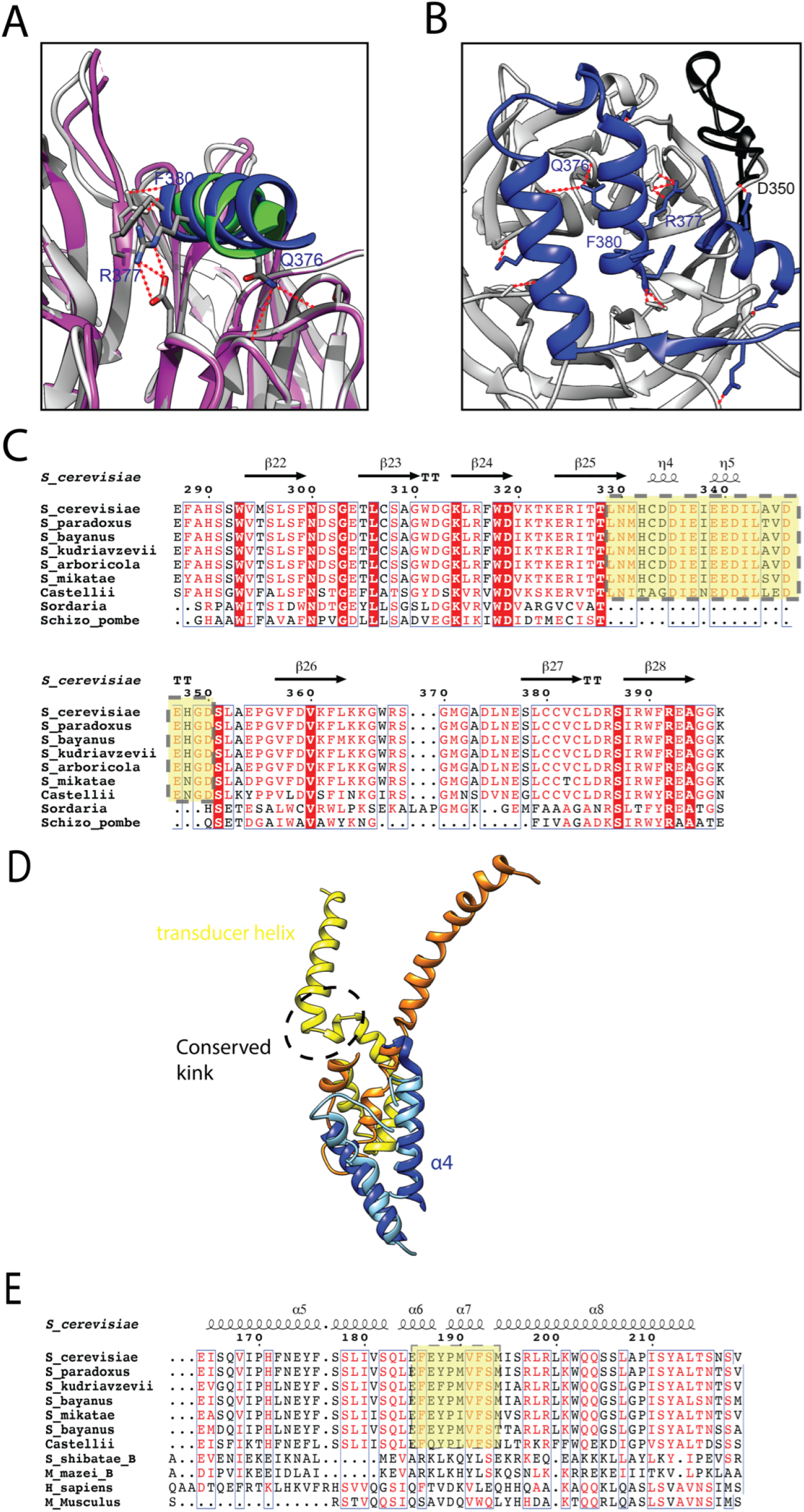
Interfaces of Spo11 with Ski8 and Rec102. (**A**) The AF model for Ski8 (gray) interaction with an alpha helix containing the conserved Spo11 QREIF_380_ motif (blue) is compared with a model (magenta and green) previously generated based on a complex of Ski8 with a similar peptide sequence in Ski3 (*25*)(*89*)). Red dashed lines indicate predicted hydrogen bonds in the RF model. (**B**) Spo11 residues 323-381 (blue) form extensive contacts with Ski8. The black segment is a part of Ski8 that appears to extend the interaction surface. Within this segment, Ski8 D350 is predicted to form a hydrogen bond with Spo11 Q323 and a salt bridge with Spo11 R320. Predicted hydrogen bonds are indicated by red dashed lines. (**C**) Sequence alignment of fungal Ski8 orthologs. The black region in panel B corresponds to an insertion (residues 329-350, highlighted in yellow) that is found in *Saccharomyces* species but not in *S. macrosporaor S. pombe*, in which Ski8 homologs also interact with Spo11. (**D**) Comparison of the Spo11–Rec102 and Top6A–Top6B interfaces. In the RF model, Spo11 residues 42 to 121 are shown in blue (a b-sheet in this region omitted for clarity) and Rec102 residues 164 to 229 are shown in yellow. From the crystal structure of the *M. mazei* Top6A–Top6B complex, Top6A residues 15 to 68 are shown in cyan and Top6B residues 440 to 508 are shown in orange. The helix labeled “a4” is the first helix in the WHD of Spo11. (**E**) Multiple sequence alignment of transducer helices from yeast Rec102, mammalian Top6BL, and archaeal Top6B proteins. The black box highlights the sequence corresponding to the predicted kink in the transducer helix of the yeast proteins, not apparent in Top6B or Top6BL.

**Figure S9.**
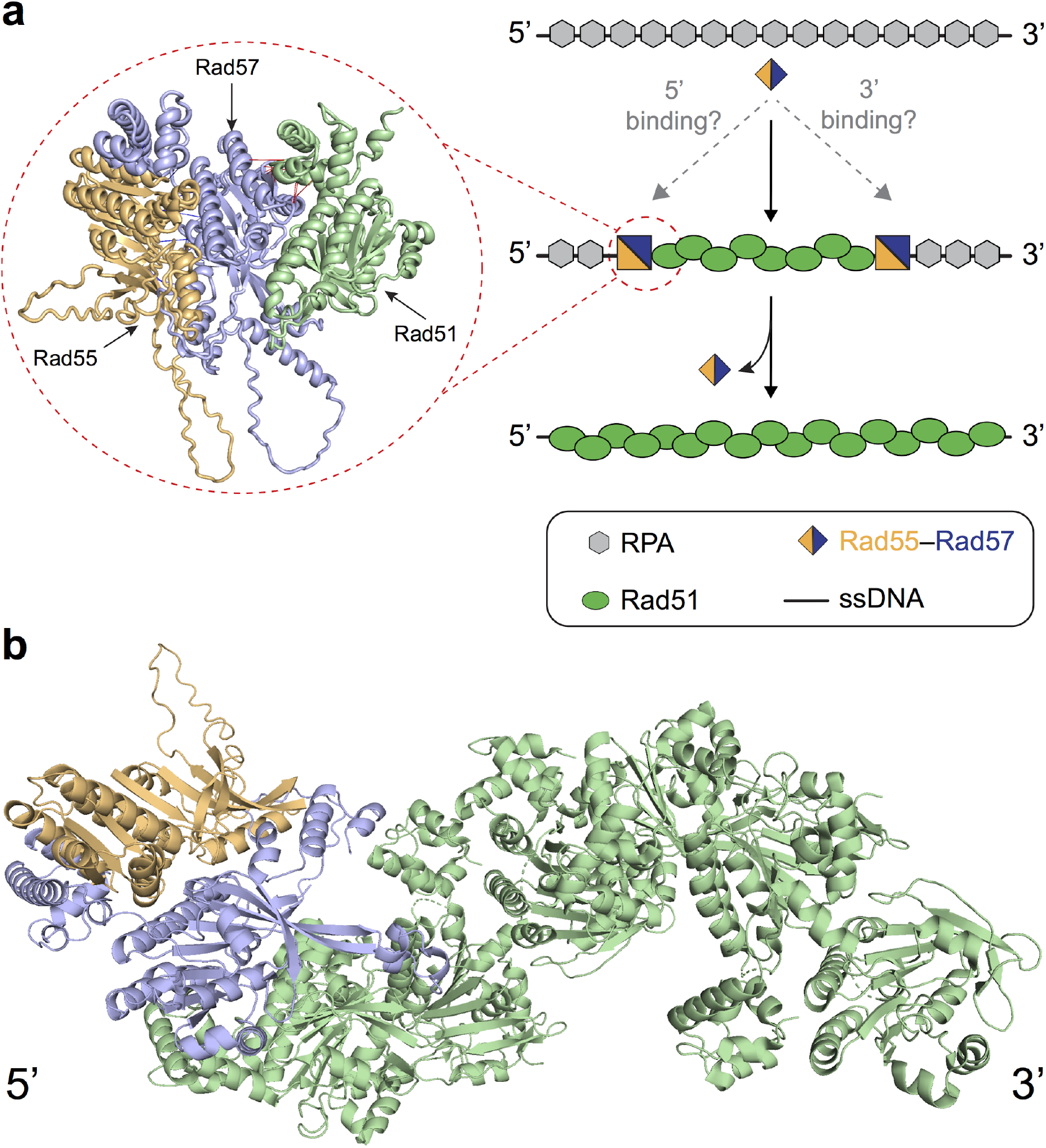
Model for Rad55–Rad57 binding to the Rad51–ssDNA filament. (**A**) Schematic for Rad55–Rad57 mediated Rad51 filament assembly during homologous recombination. Rad51 recombinase must displace replication protein A (RPA) bound to single–stranded DNA (ssDNA) to form Rad51–ssDNA filaments that carry out DNA recombination. Rad55–Rad57 acts as a chaperone, facilitating a faster and more extensive displacement of RPA by Rad51. The specific polarity of interaction between Rad55–Rad57 and the Rad51–ssDNA filament is unknown, but is expected to influence whether filament growth is stimulated in the 5’→3’ or 3’→5’ direction. Our model suggests that Rad55–Rad57 may be transiently binding at the 5’ end of a Rad51–ssDNA filament through an interaction between Rad57 and Rad51 (inset). (**B**) Putative model for Rad55–Rad57 binding at the 5’ end of a Rad51 filament.

**Figure S10.**
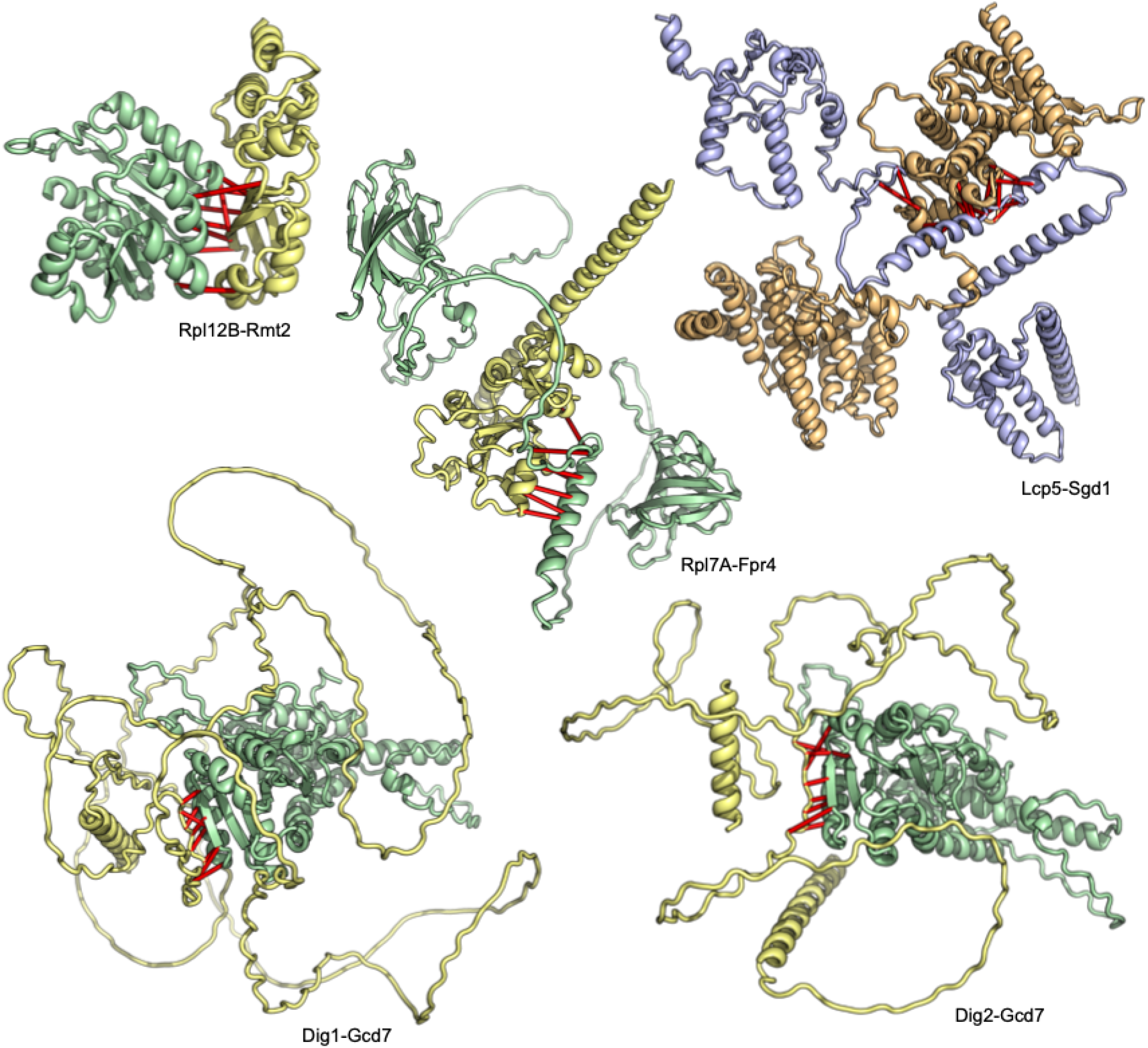
Structures of complexes involved in translation and ribosome regulation. Top predicted residue-residue contacts are indicated with bars. Pair color indicates the method of identification from Fig. 1D; experiment-guided pairs are yellow and green and “*de novo”* pairs are blue and light-orange.

**Figure S11.**
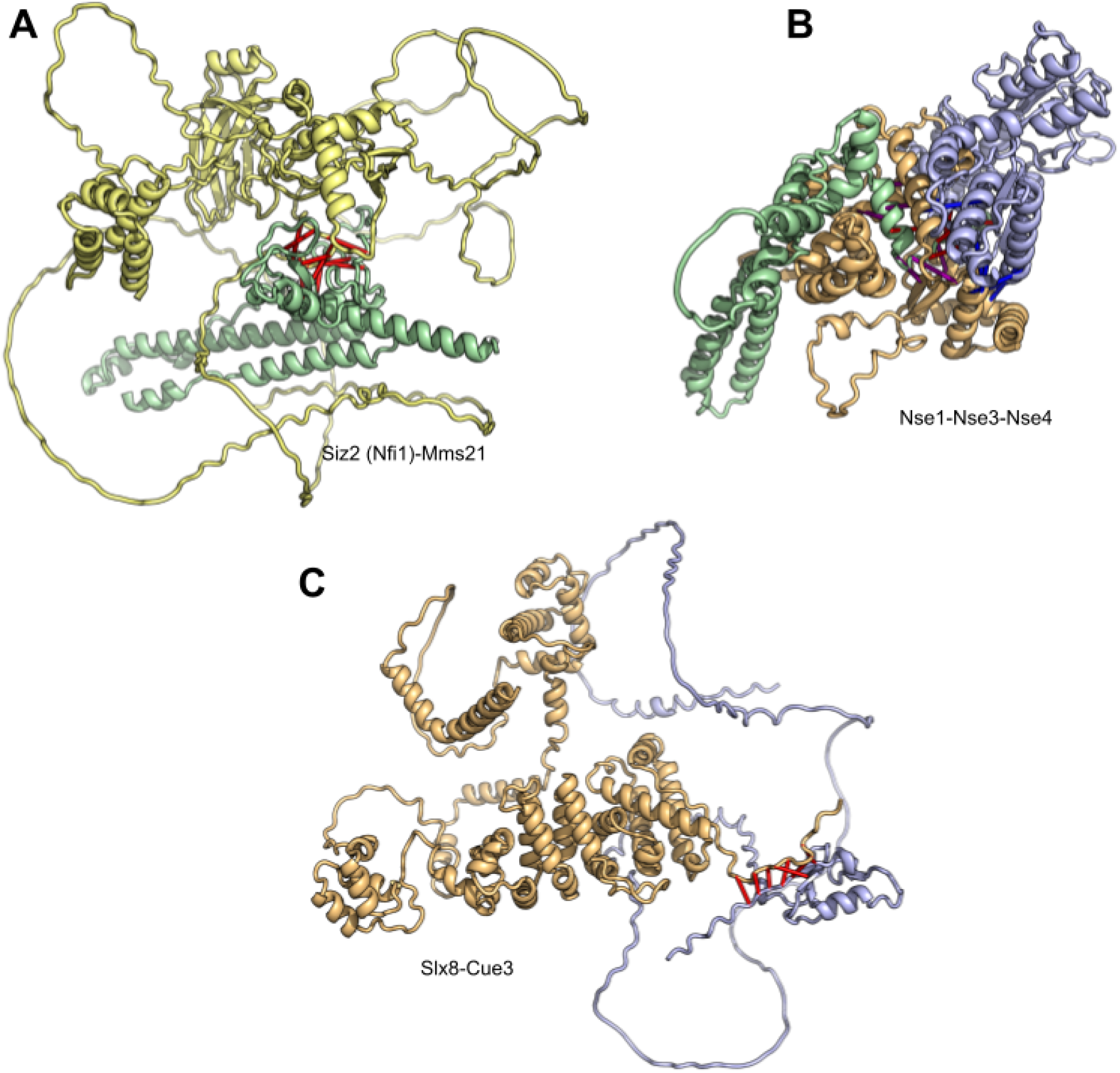
Complexes involving SUMO and ubiquitin ligases. Top predicted residue-residue contacts are indicated with bars. Pair color indicates the method of identification from Fig. 1D and Fig. 5; experiment-guided pairs are yellow and green (panel A) and “*de novo”* pairs are blue and light-orange (panel C).

**Figure S12.**
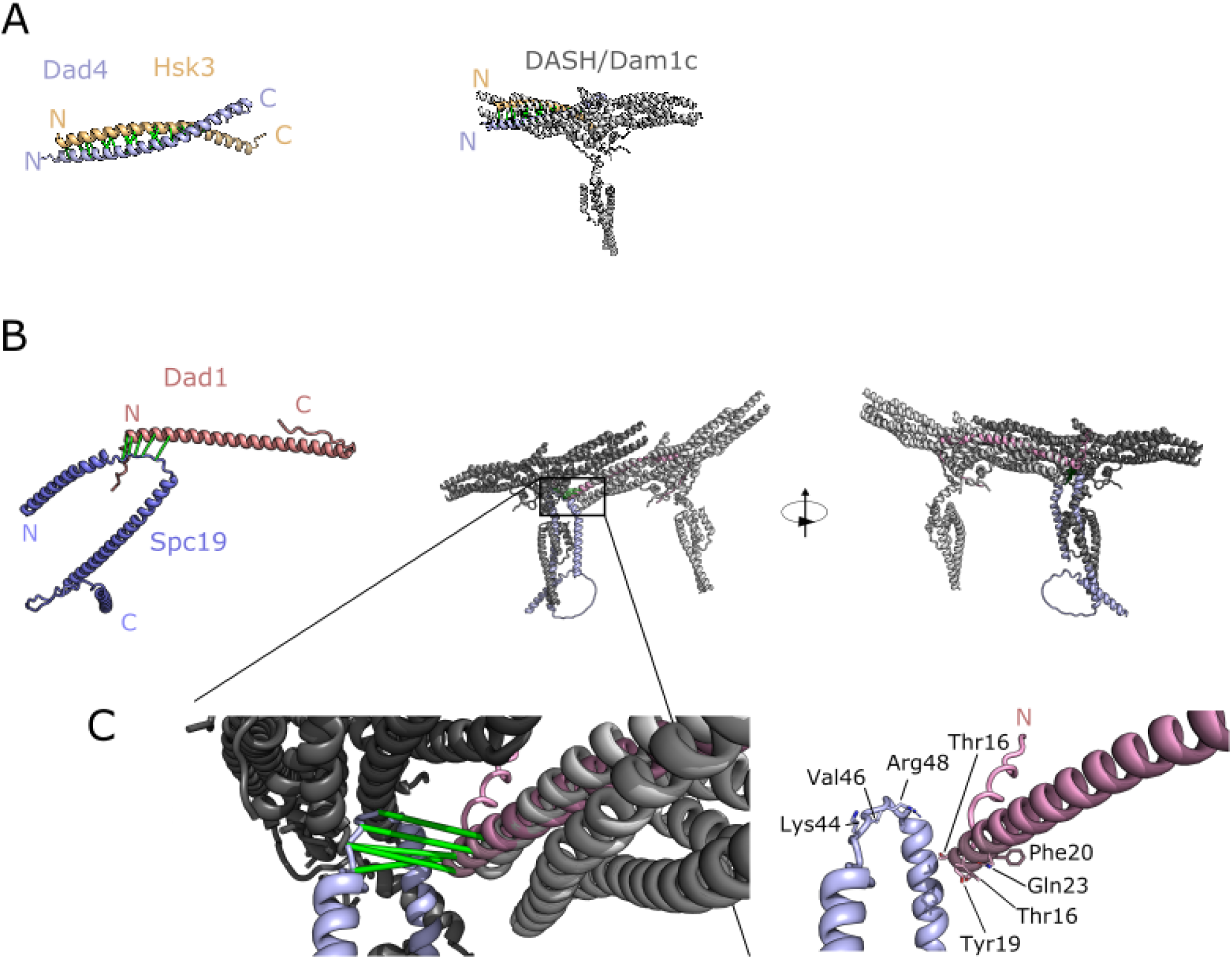
Predicted inter and intra-decamer interactions in the DASH/Dam1 complex. **(A)** Left: Predicted dimer between Dad4 (blue) and Hsk3 (gold) proteins, with predicted contacts shown in green. Right: The Dad4-Hsk3 dimer aligned with the structure of the DASH/Dam1 decamer complex from *C. thermophilus* (grey;PDB:6CFZ; (*59*)). (**B**) Left: Predicted dimer between Spc19 (blue) and Dad1 (pink) with predicted contacts shown in green. Center & right: The Spc19 Dad1 dimer aligned with the structure of the DASH/Dam1 decamer complex from *C. thermophilus* (grey;PDB:6CFZ; (*59*)) in the context of ring formation. The image on the right has been rotated 180° about the Y axis. (**C**) A zoomed view of the Dad1-Spc19 interactions with (left) and without (right) the *C. thermophilus* structure visible.

**Figure S13.**
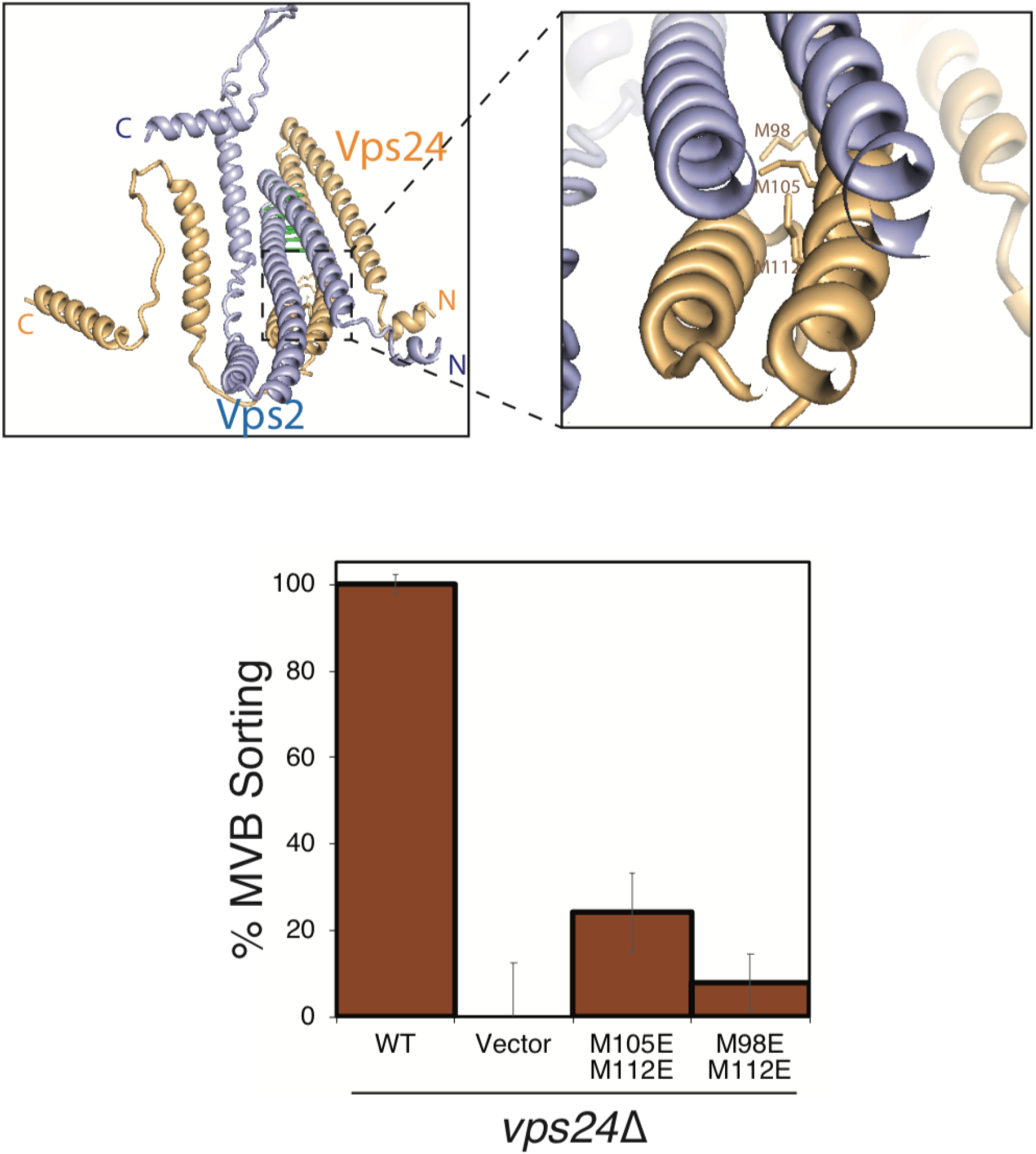
Predicted Vps2-Vps24 complex structure is consistent with unpublished mutagenesis data. Predicted ESCRT-III interface mutations inhibit cargo sorting functions. (**A**) Predicted structure of the yeast ESCRT-III complex Vps24-Vps2. Figure on the right represents a zoomed-in image of the box with dotted-lines. Residues M98, M105 and M112 in the helix-3 region of Vps24 are highlighted in “sticks” representation. (**B**) Data represent flow-cytometry cargo sorting assay in *S. cerevisiae.* The methionine transporter Mup1 tagged with pHluorin (Mup1-pHluorin) was used as cargo. Upon sorting to the vacuole, the fluorescence of pHluorin is quenched, which is quantified as 100% for WT and 0% for the empty vector. Error bars represent standard deviation of three independent experiments.

**Figure S14.**
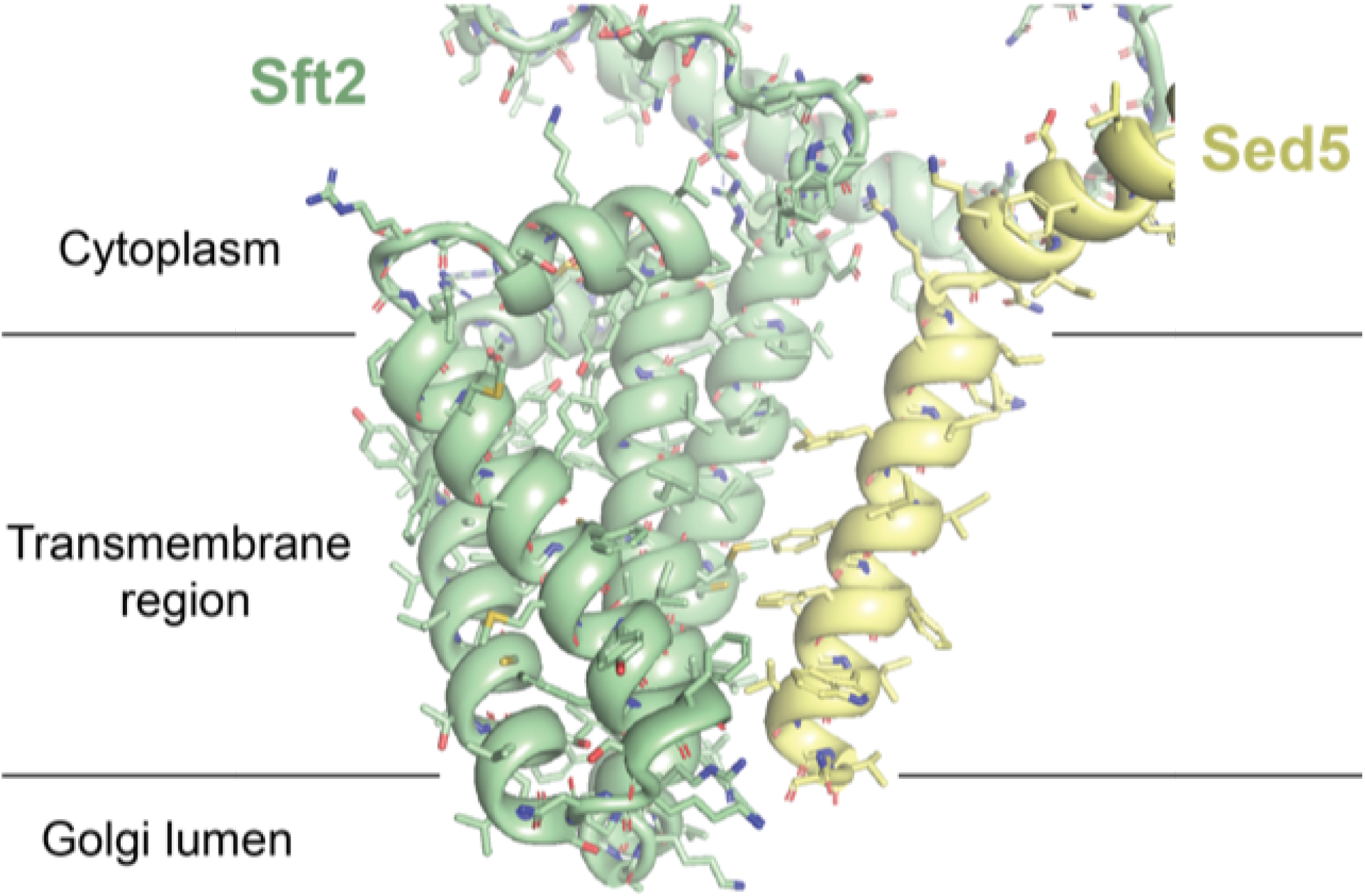
Sft2-Sed5 annotated interface. Close-up view of the predicted structure of the Sft2-Sed5 interface. Sft2 has been implicated as a regulator of SNARE function in membrane fusion. The transmembrane helix of the SNARE Sed5 (yellow) is predicted to interact with two transmembrane helices of Sft2 (green). The expected transmembrane domain boundaries are depicted as thin black lines.

**Figure S15.**
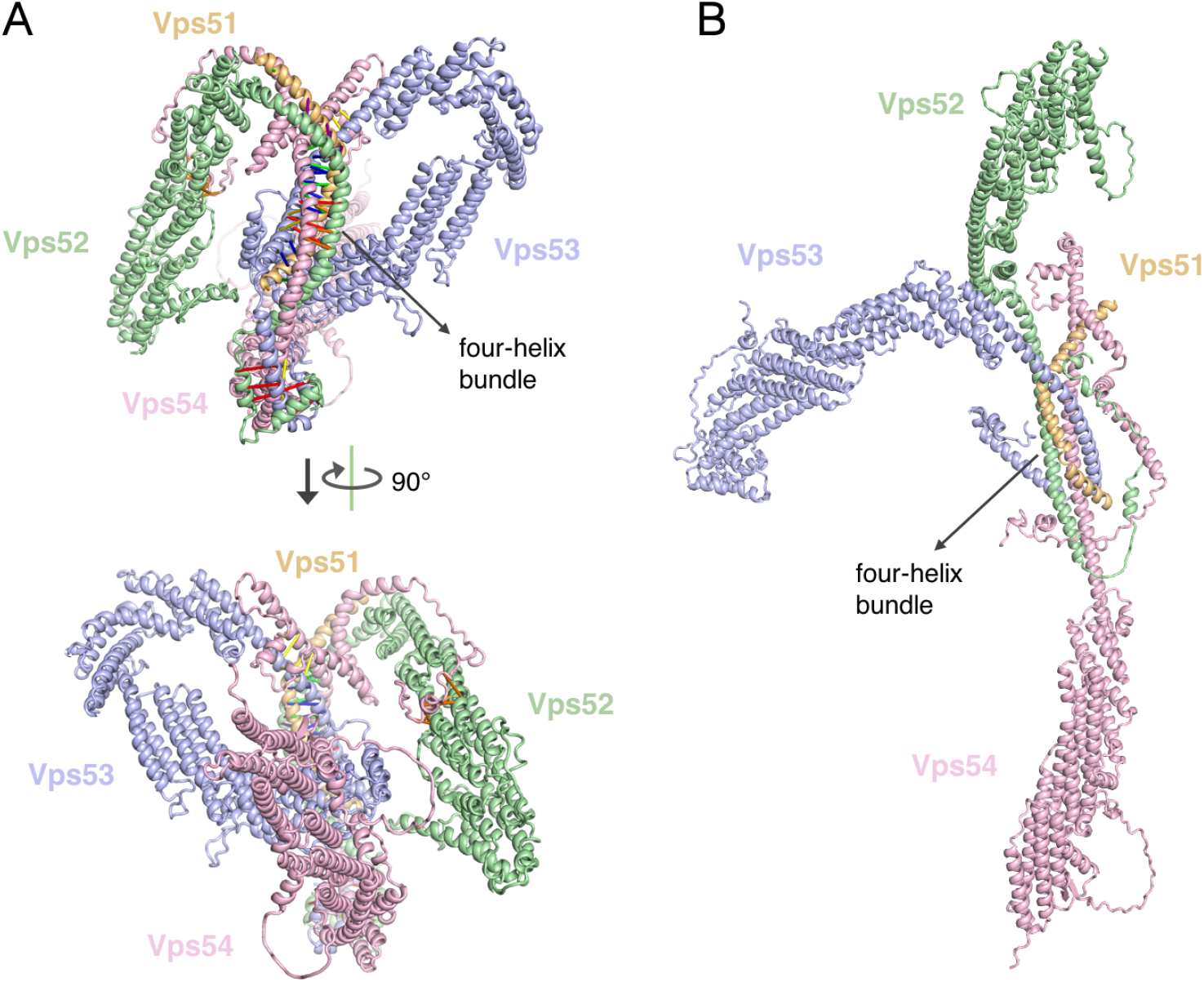
Predicted GARP complex structure is consistent with negative stain class averages. (**A**) Predicted model of the GARP complex is shown in two orientations. Predicted residue-residue contacts are indicated with bars. (**B**) Model constructed by superimposing individual AF2 predictions for Vps52, Vps53, and Vps54 onto the central four-helix bundle. The resulting model shows a different overall architecture.

**Figure S16.**
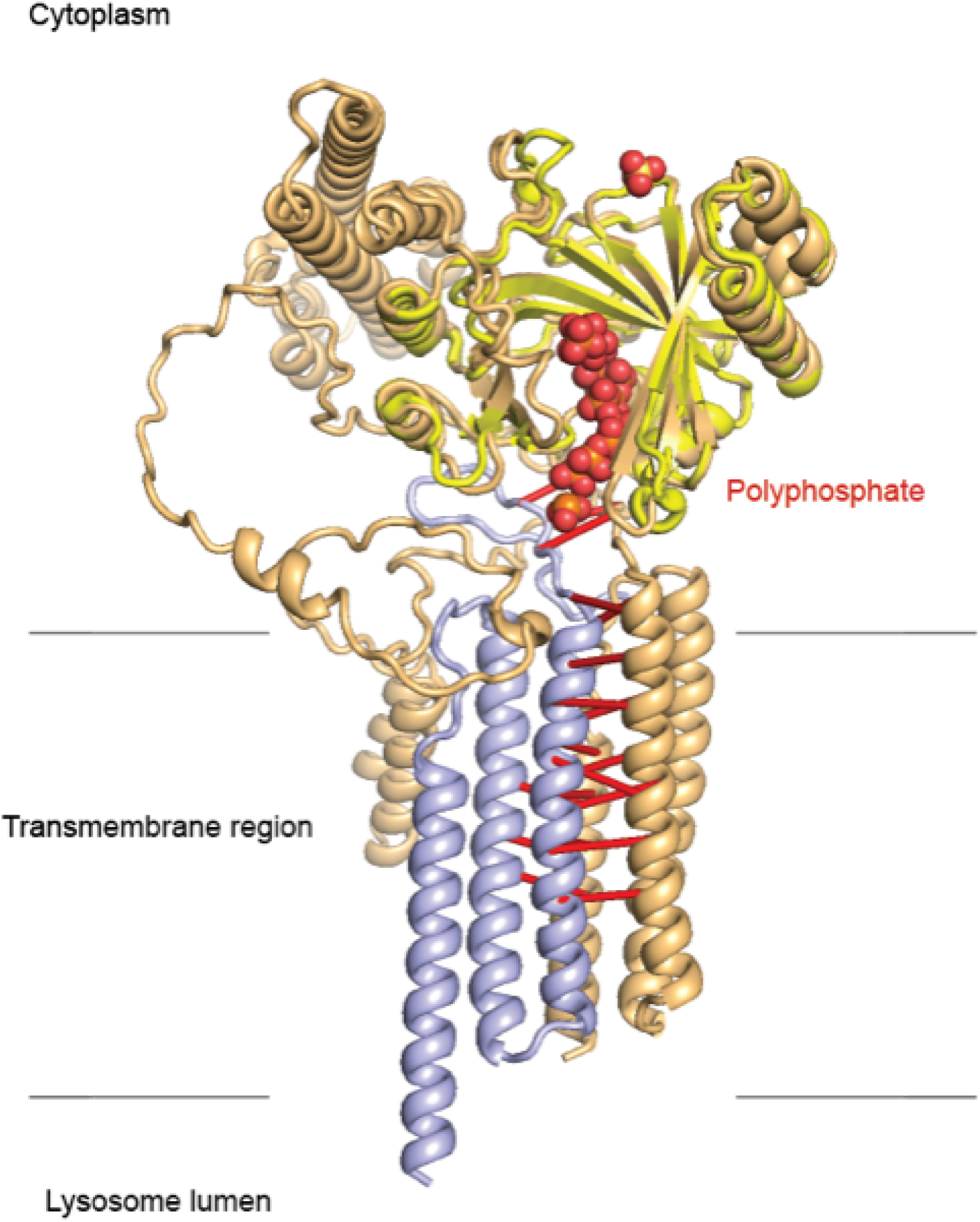
Structural model for subunits of the VTC complex. The VTC complex synthesizes polyphosphate and transports it into the vacuole/lysosome lumen. This model was produced by superposition of the crystal structure (PDB: 3G3Q, Chain A) of the VTC active site from Vtc4 (bright yellow) with bound polyphosphate (red) onto our predicted structure of the Vtc1-Vtc4 complex (colored lightblue and lightorange, respectively). The expected transmembrane domain boundaries are depicted as thin black lines.

**Figure S17.**
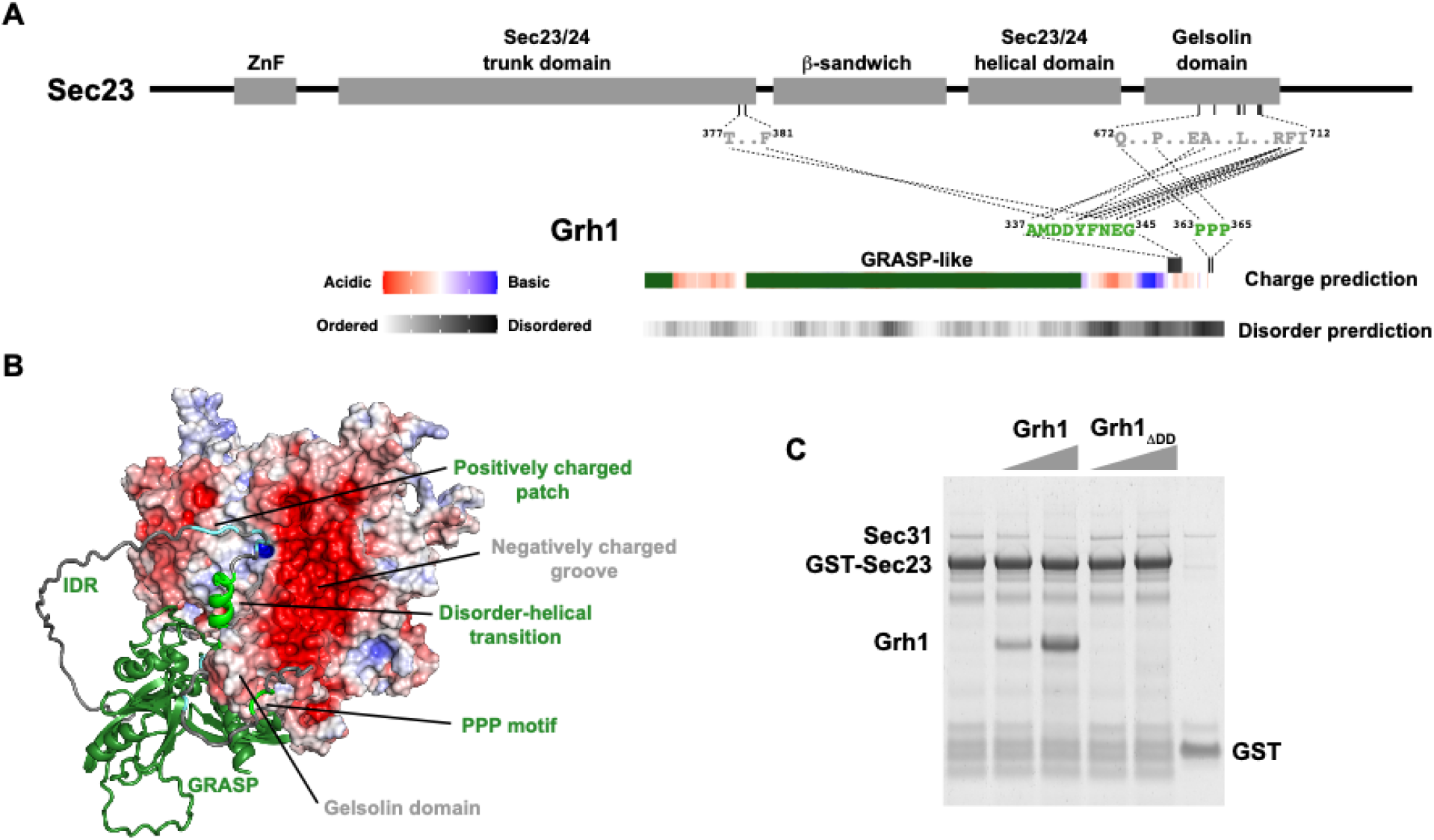
Predicted multivalent complex between Grh1-Sec23. (**A**) Domains and predicted interactions of Sec23 (top) and Grh1 (bottom). Grh1 contains a C-terminal IDR which hosts a positively charged cluster and two regions that are predicted to interact with the gelsolin domain in Sec23 (dashed lines). (**B**) Predicted structure of the Sec23-Grh1 complex displayed two interfaces between the IDR (grey) in Grh1 and Sec23 (rendered by surface charge): a helical motif involving disorder-to-helical transition (light green) and a polyproline PPP motif (dark green). The flanking positively charged patch in the IDR may sample the negatively charged groove in Sec23 to anchor the multivalent interface. (**C**) GST pulldown assay illustrating the importance of interaction between C-terminal IDR of Grh1 and Sec23. Full length Grh1 was able to compete with outer coat protein Sec31 upon recruitment by Sec23; deletion of the C-terminal IDR which harbours the predicted multivalent interface completely abolished the recruitment of Grh1 to Sec23.

**Figure S18.**
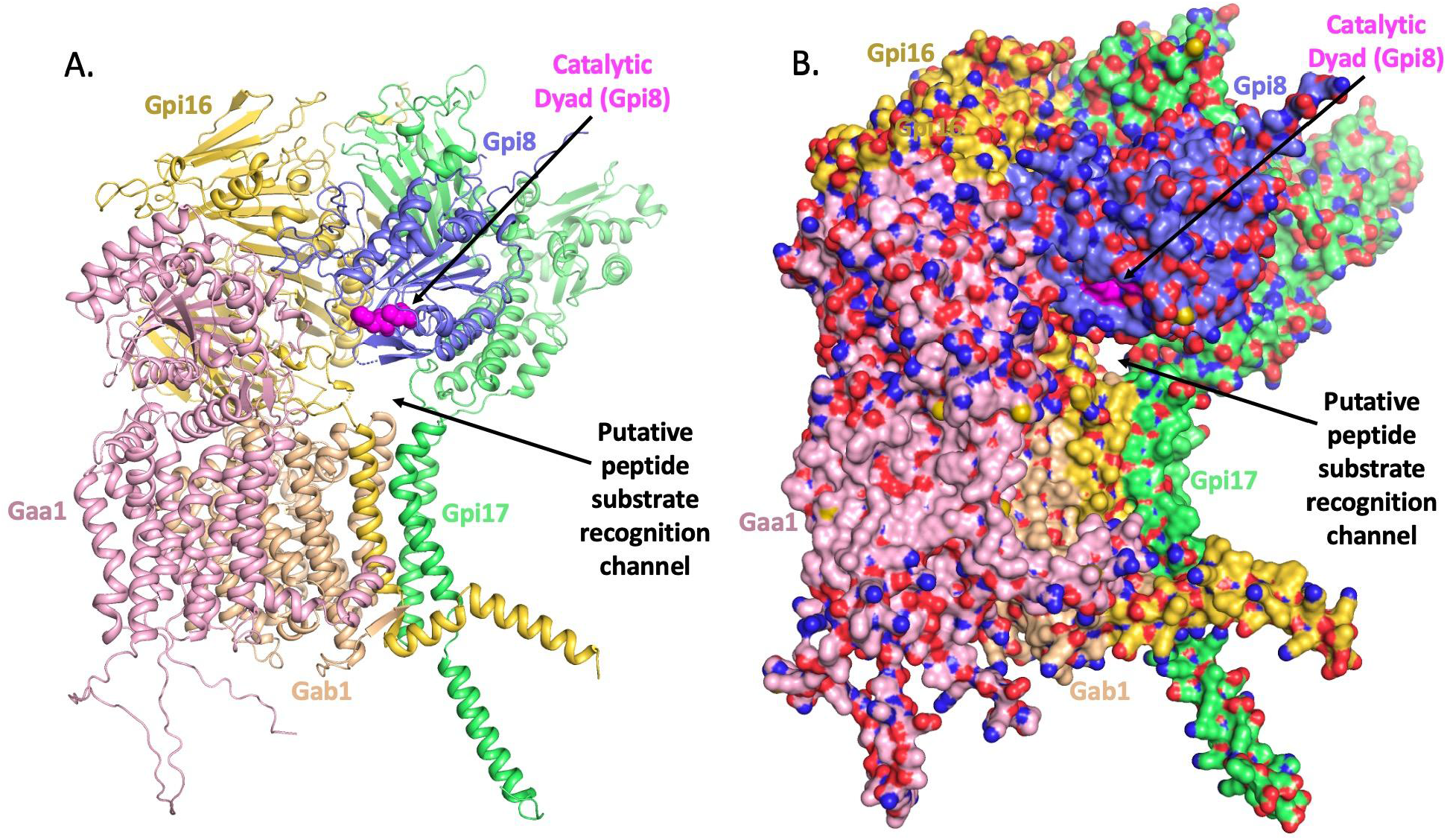
C-terminal GPI-T signal sequence recognition tunnel suggested by GPI-T complex model. GPI-T recognizes a C-terminal signal sequence composed of a pattern of small, hydrophilic, and then hydrophobic residues. The model of GPI-T reveals a putative channel between the Gpi8, Gpi16, and Gpi17 subunits. The position of this tunnel is shown in a cartoon model (**A**) and in a surface model (**B**) of GPI-T. The side chains of Ala168 in Gpi8 and Phe500 in Gaa1 were hidden for better visualization of this channel. The N-terminal signal peptides were removed from Gpi8 (residues 1-22) and Gpi16 (residues 1-19). For Gpi8, only residues 23-306 are included in the model. Each subunit is color coded as indicated, with the catalytic dyad in Gpi8 (Cys199 and His157) highlighted in magenta.

